# Cell type-selective secretome profiling in vivo

**DOI:** 10.1101/2020.09.18.303909

**Authors:** Wei Wei, Nicholas M. Riley, Andrew C. Yang, Joon T. Kim, Stephanie M. Terrell, Veronica L. Li, Marta Garcia-Contreras, Carolyn R. Bertozzi, Jonathan Z. Long

## Abstract

Secreted polypeptides are a fundamental biochemical axis of intercellular and endocrine communication. However, a global understanding of composition and dynamics of cellular secretomes in intact mammalian organisms has been lacking. Here, we introduce a proximity biotinylation strategy that enables labeling, detection, and enrichment of secreted polypeptides in a cell type-selective manner in mice. We generate a proteomic atlas of hepatocyte, myocyte, pericyte, and myeloid cell secretomes by direct purification of biotinylated secreted polypeptides from blood. Our secretome atlas validates known cell type-protein pairs, reveals secreted polypeptides that distinguish between cell types, and identifies new cellular sources for classical plasma proteins. Lastly, we uncover a dynamic and previously undescribed nutrient-dependent reprogramming of the hepatocyte secretome characterized by increased unconventional secretion of the cytosolic enzyme BHMT. This secretome profiling strategy enables dynamic and cell-type dissection of the plasma proteome and the secreted polypeptides that mediate intercellular signaling.

## MAIN

Secreted polypeptides are a fundamental biochemical axis of intercellular and endocrine communication. In mammals, these extracellular molecules are present in blood plasma, can be dynamically regulated by physiological perturbations, and regulate critical homeostatic processes. For instance, pancreatic beta cells secrete insulin in response to increased blood glucose^1^ and endothelial cells secrete chemokines and extracellular matrix components in response to local, microenvironmental changes^2^. However, a global understanding of cellular secretomes and their dynamic regulation, especially within the context of an intact mammalian organism, has been lacking. Shotgun proteomics of conditioned media from cultured cells does not necessarily capture the physiologic secretome in vivo^3^. Genomic predictions based on the presence of an N-terminal signal peptide ignores the diversity of alternative secretion pathways (e.g., unconventional secretion, ectodomain shedding)^4^. Lastly, none of these approaches provide temporal information on dynamic secretion events regulated by complex physiological perturbations such as nutrient availability or disease states.

Recently, secretome profiling technologies based on bio-orthogonal tagging of secreted polypeptides have emerged^5–9^. By metabolic incorporation of analytical handles for downstream detection and enrichment of secreted polypeptides, these approaches have provided a chemical strategy for potentially overcoming the limitations associated with more classical secretome profiling approaches. For instance, addition of azidohomoalanine, a methionine analog, to serum-containing media leads to azidoylation of secreted proteins in cell culture^5^. An engineered tRNA synthetase that incorporates azido-phenylalanine has also been used to label secreted proteins from tumor xenographs^5–7^. However, none of these strategies have yet been successfully translated to label endogenous cellular secretomes in mice to date, likely due to inefficient in vivo metabolic incorporation or other untoward effects associated with the high levels of unnatural amino acid administration.

To complement all of these approaches, here we introduce a proximity biotinylation strategy applicable for cell type-selective secretome profiling in mice. Our approach is based on in vivo biotinylation of intracellular proteins and then following those biotinylated proteins as they are released extracellularly. By expressing these constructs in a cell type-selective manner, we profile in vivo cellular secretomes by direct enrichment of biotinylated proteins from blood plasma. Our in vivo atlas of four cell type secretomes (hepatocytes, myocytes, pericytes, and myeloid cells) validates known cell type-protein pairs, reveals secreted proteins that distinguish between cell types, and identifies new cellular sources for classical plasma proteins. Lastly, we uncover a previously undescribed fructose-dependent dynamic reprogramming of hepatocyte secretion characterized by increased selective unconventional protein export of the cytosolic enzyme BHMT (betaine-homocysteine S-methyltransferase). This secretome profiling strategy therefore enables cell-type specific dissection of secreted plasma proteins and their dynamic regulation by physiological perturbations.

## RESULTS

### Biotinylation of secreted polypeptides in cell culture

Unlike many of the previous efforts that have relied on metabolic incorporation of bio-orthogonal reagents, we reasoned that proximity labeling^10–12^ might be a suitable alternative method for biotinylation of intracellular and secreted polypeptides (**Figure 1a**). As our labeling reagent, we used the recently engineered TurboID owing to its rapid kinetics and robust activity in oxidizing environments^12, 13^. Because of the reported compartment-specific nature of proximity labeling, we initially generated three different constructs targeted to distinct subcellular compartments. We first constructed a luminally-oriented, membrane-tethered TurboID construct by appending an N-terminal signal peptide and C-terminal transmembrane domain from *Pdgfrb*^14^ (Mem-TurboID, **Supplemental Figure 1a**). In parallel, we also used cytoplasmic and luminally-oriented ER-localized TurboID constructs (Cyto-TurboID and ER-TurboID, respectively)^12^ which had been previously described for intracellular labeling but had not been evaluated for labeling secreted polypeptides (**Supplemental Figure 1a**). We anticipated that both Mem-TurboID and ER-TurboID would predominantly capture classical secretion events which occur by ER/Golgi trafficking, while the Cyto-TurboID construct might be more suited for proteins that undergo unconventional protein export (**Supplemental Figure 1b**). The subcellular localization for each construct determined by immunofluorescence microscopy. As expected, robust vesicular and plasma membrane signal was observed for Mem-TurboID (**Supplemental Figure 1c**). Cyto-TurboID was observed to be diffusely distributed across the cytoplasm and ER-TurboID staining was restricted to a tubular perinuclear network (**Supplemental Figure 1c**).

**Figure 1.**
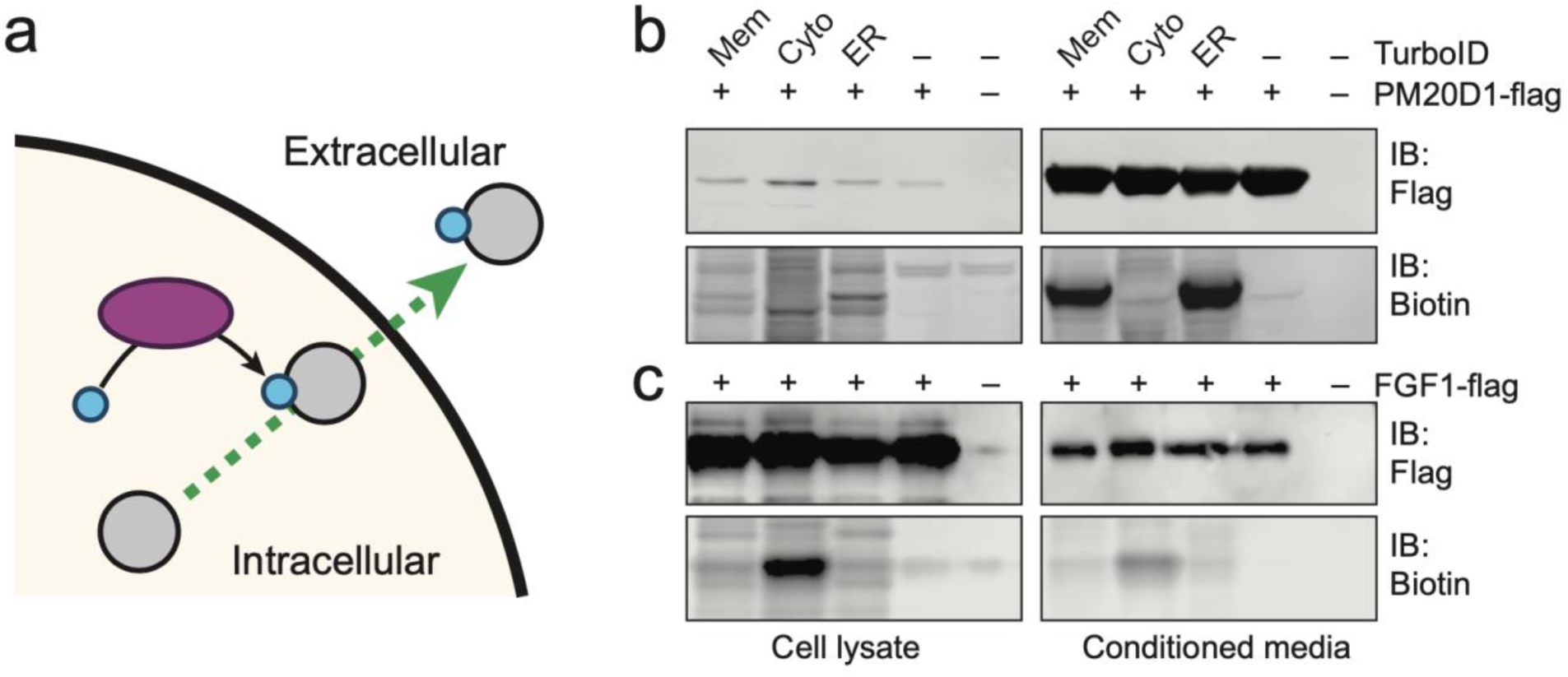
Biotinylation of secreted proteins in cell culture. **(a)** Schematic of the proximity labeling strategy used for tagging secreted polypeptides. Grey circle represents secreted proteins, blue dot indicates biotinylation, and purple oval represents proximity labeling reagent. **(b, c)** Anti-flag or streptavidin blotting of conditioned media and cell lysates from HEK293T cells transfected with the indicated proximity labeling constructs and the classically secreted PM20D1-flag **(b)** or the unconventionally secreted FGF1-flag **(c)**. Proximity labeling was initiated one day post transfection by switching cells into serum-free media in the presence of 500 µM biotin for 18 h. Experiments were performed three times and similar results were obtained.

We next characterized biotinylation of secreted proteins in cell culture. Each proximity labeling construct was co-transfected with a classically secreted protein (PM20D1^15, 16^) or an unconventionally secreted protein (FGF1^17^). Biotinylation was initiated on the day following transfection (500 µM biotin in serum free media) and conditioned media was harvested 18 h later. Extracellular PM20D1 was robustly biotinylated by the two lumenally-oriented constructs Mem-TurboID and ER-TurboID, but not by Cyto-TurboID (**Figure 1b**). No biotinylation was observed when PM20D1 was transfected alone, thereby confirming the specificity of the labeling event. By contrast FGF1, was robustly biotinylated by Cyto-TurboID, and to a lesser extent by the two other luminally-oriented constructs (**Figure 1c**). Based on quantitation of fractional biotinylation of flag-tagged proteins in the conditioned media, we estimate a labeling efficiency of 7%, 38%, and 22% for PM-, Cyto-, and ER-TurboID, respectively (**Supplemental Figure 1d**). We therefore conclude that this suite of three proximity labeling constructs can biotinylate a diversity of polypeptide secretion events in cell culture, with some apparent cross-labeling of secretory pathways. These in vitro data also demonstrate that using all three constructs may be useful for capturing a broader swath of secretion events compared to a single construct alone.

### In vivo labeling of the hepatocyte secretome

To determine whether proximity biotinylation would be feasible for labeling endogenous cellular secretomes in mice, we cloned each of the constructs into adeno-associated virus (AAV) plasmids with the hepatocyte-directed thyroxine binding protein (*Tbg*) promoter^18^ (**Figure 2a**). We selected hepatocytes as an initial cell type because of their robust secretory capacity and ease of genetic manipulation. Following tail vein transduction of AAVs, Western blotting using an anti-V5 antibody to detect TurboID protein demonstrated that each AAV-*Tbg* virus exhibited the expected liver-restricted expression (**Figure 2b**). Immunofluorescence analysis of frozen liver sections revealed that >95% of hepatocytes were transduced in virus-injected versus control mice, though variation in expression levels between hepatocytes was also observed (**Figure 2c**). ALT was not elevated in transduced versus control animals (**Supplementary Figure 2**), demonstrating that the doses of viruses used here do not cause overt organ damage.

**Figure 2.**
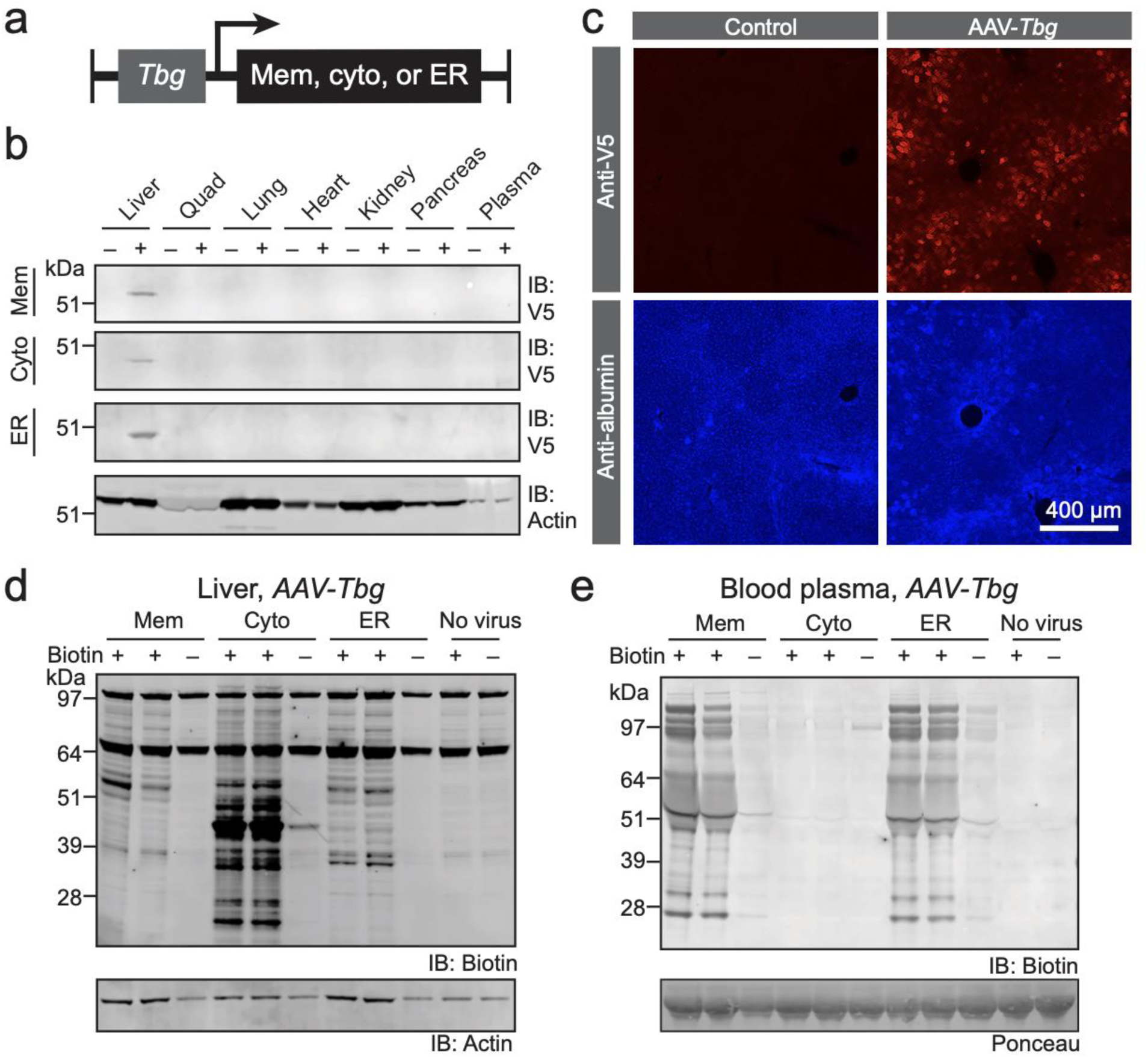
Biotinylation of hepatocyte secretomes in vivo. **(a)** Cartoon schematic of the adeno-associated virus constructs driven by the hepatocyte-specific *Tbg* promoter. **(b)** Anti-V5 blotting of a panel of murine tissues following transduction by AAV-*Tbg* viruses. (+) indicates AAV transduction and (–) indicates no viral transduction. **(c)** Anti-V5 (top panels) or anti-albumin (bottom panels) immunofluorescence of frozen liver sections from AAV-*Tbg*-ER or control mice. **(d,e)** Streptavidin blotting (top panels) or loading controls (bottom panels) of liver lysates **(d)** or blood plasma **(e)** from mice transduced with the indicated AAV and then treated with vehicle (–) or biotin (24 mg/kg/day, intraperitoneally, for three consecutive days). Tissues were harvested 24 h after the final biotin injection. For AAV-*Tbg* virus transduction, male C57BL/6J mice were 6-8 weeks old and transduced for 1 week prior to in vivo biotin labeling.

Next, we evaluated tissue biotinylation by injecting mice with biotin (24 mg/kg/day for three consecutive days, intraperitoneal injection)^19^. Liver was harvested one day after the final injection. As expected, biotinylation proteins were detected in livers from AAV-*Tbg* transduced mice (**Figure 2d**). Importantly, control experiments omitting either AAV transduction or biotin administration revealed minimal signal over background (**Figure 2d**), establishing the virus- and biotin-dependence of the observed labeling events. Lastly, to determine whether secreted proteins from hepatocytes were also biotinylated, we analyzed blood plasma from these same animals. As shown in **Figure 2e**, robust virus- and biotin-dependent appearance of biotinylated proteins in blood plasma was over background. Both AAV-*Tbg*-Mem and AAV-*Tbg*-ER viruses gave robust blood plasma biotinylation signals that were largely overlapping in pattern. By contrast, the AAV-*Tbg*-Cyto virus showed less biotinylated protein in the blood over background. We therefore conclude that these AAV-*Tbg* viruses enable robust biotinylation of the hepatocyte secretome in vivo.

To determine whether additional biotin administration methods could also efficiently label secretomes in vivo, biotin (0.5 mg/ml) was provided in drinking water to AAV-*Tbg*-ER transduced mice. Fluid intake was equivalent between biotin-supplemented versus standard control water (**Supplemental Figure 3a**), establishing that this dose of biotin is well-tolerated. After a three-day labeling period, plasma was harvested and analyzed by streptavidin blotting. Biotinylated hepatocyte-derived plasma proteins were once again observed (**Supplemental Figure 3b,c**). By comparison of biotinylated proteins in blood plasma, we estimate an increased ∼6-fold labeling by biotin water versus intraperitoneal injection. These data establishing that intraperitoneal and biotin-supplemented water administration methods are both feasible for in vivo secretome labeling, with biotin-supplemented water providing more efficient in vivo labeling than intraperitoneal delivery.

### Proteomic characterization of the hepatocyte-secreted plasma proteins

We next performed a shotgun proteomics experiment to determine the identities of hepatocyte-derived polypeptides secreted into the blood (**Supplemental Table 1**). Towards this end, biotinylated proteins from blood plasma of virus-transduced mice was purified on streptavidin beads, digested using an S-trap protocol, and analyzed by liquid chromatography tandem-mass spectrometry (LC-MS/MS). As an additional control for intracellular labeling, we also performed shotgun proteomics of intracellular biotinylated proteins immunoaffinity purified from liver lysates of virus-transduced mice (**Supplemental Table 2**). In the liver, we detected 31,003 unique peptides from 218,304 peptide spectral matches corresponding to 2,994 proteins, establishing that the liver proteome is broadly biotinylated. In blood plasma, 4,779 unique peptides were detected from 77,216 peptide spectral matches, corresponding to 303 proteins.

Principal component analysis showed clear separation of each of biotinylated plasma proteins from each of the AAV-*Tbg* viruses versus control mice (**Figure 3a**). In total, 56, 8, and 65 proteins were found to be statistically significantly enriched in each pair-wise comparison of AAV-*Tbg*-Mem, AAV-*Tbg*-Cyto, or AAV-*Tbg*-ER versus control (**Supplemental Figure 4**). To quantitatively understand the similarities and differences between these three sets of proteins in an unbiased manner, we used hierarchical clustering to categorize significantly distinct proteins as determined by ANOVA (**Figure 3b**). The largest cluster of 56 plasma proteins (highlighted in teal color, **Figure 3b**) was commonly enriched in either AAV-*Tbg*-Mem or AAV-*Tbg*-ER viruses versus control, but not AAV-*Tbg*-Cyto versus control. Gene ontology analysis for this cluster established an enrichment of extracellular signal peptide-containing secreted glycoproteins across multiple biological pathways, including protease inhibition, complement pathway, and immunity (**Figure 3c**). Manual inspection of these proteins identified many known classically secreted liver-derived plasma proteins including known lipoprotein components (e.g., APOB, APOA1, LCAT1), complement proteins (e.g., C5, C9, CFB, C4B), coagulation factors (e.g., F2, F10, F12), serine protease inhibitors (e.g., SERPIND1), and hormones (e.g., FETUB, GPLD1) (**Supplemental Table 1**). By comparison, biotinylated hepatocyte-secreted proteins labeled by AAV-*Tbg*-Cyto (highlighted in orange, **Figure 3b**) included intracellular proteins (e.g., TENS4) as well as additional classically secreted proteins not labeled by the other two constructs (e.g., FIBA, PLMN).

**Figure 3.**
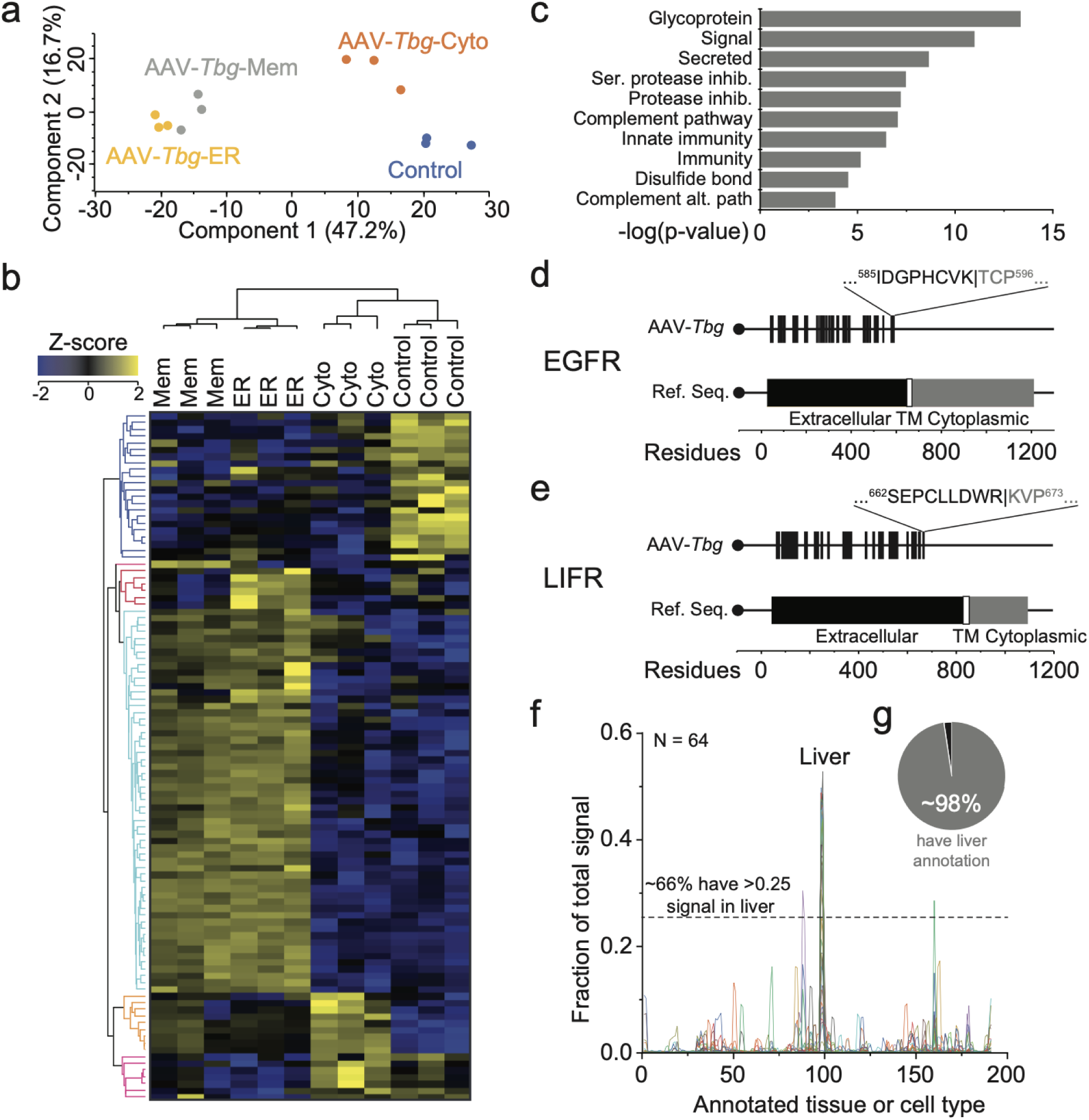
Proteomic characterization of the hepatocyte secretome. **(a)** Principal component analysis of streptavidin-purified plasma proteins from mice transduced with the indicated AAV-*Tbg* for one week and then injected with biotin (24 mg/kg/day, intraperitoneally, for three consecutive days). N = 3 mice/group. **(b)** Hierarchical clustering by Z-score intensities of all differentially detected streptavidin-purified plasma proteins from AAV-*Tbg*-Mem, AAV-*Tbg*-Cyto, AAV-*Tbg*-ER, or control mice. **(c)** Gene ontology analysis of the cluster highlighted in teal. **(d,e)** Schematic of detected peptides for the transmembrane receptors EGFR **(d)** and LIFR **(e)** mapped onto their respective reference sequences with annotated domains indicated below. Observed cleavage sites are indicated by “|” in the amino acid sequence above each protein map, with the black text showing residues from the most C-terminal peptide detected. **(f)** Relative expression of significantly enriched proteins in AAV-*Tbg* streptavidin-purified plasma across 191 different tissues and cell types available in BioGPS. Each line represents one protein. The dashed horizontal line indicates which proteins had at least one fourth of their total signal in a given tissue. **(g)** Pie graph showing percentage of enriched proteins with liver expression, calculated by comparing liver expression to the median expression value across all tissues. Proteins with liver expression greater than the median value were considered as “liver proteins.”

Biotinylated transmembrane receptors were also detected in our AAV-*Tbg* plasma proteomic datasets (**Figure 3d,e**). Consistent with detection of ectodomain shedding from hepatocytes, the tryptic peptides detected for each EGFR and LIFR mapped exclusively to the annotated extracellular domains. Furthermore, we could estimate the site of cleavage based on detection of the most C-terminal peptide detected (**Figure 3d,e**). In fact, our predicted ectodomain cleavage site for EGFR resides in domain IV, the analogous binding region for the anti-neoplastic antibody Trastuzumab that prevents EGFR shedding^20^. These data demonstrate that ectodomain shedding events can be directly mapped using this approach and establish that hepatocytes are sources of circulating soluble EGFR and LIFR in blood plasma.

Lastly, we performed additional analyses to determine the cell type specificity of our genetic strategy. All proteins detected and enriched in AAV-*Tbg*-Mem and AAV-*Tbg*-ER viruses were pooled and their tissue expression collected using BioGPS as a reference database. Expression profiles were collected across 191 tissues and cell lines in BioGPS^21^ and a “normalized” mRNA expression profile was determined for each biotinylated plasma protein (see **Methods**). While a subset of the biotinylated plasma proteins exhibited diverse expression across tissues as indicated by the distribution of peaks, liver emerged as the predominant tissue with the highest representation (**Figure 3f**). In fact, 66% (42 out of 64) of proteins had at least one quarter of their total expression (a signal of 0.25) in liver (**Figure 3f**, dashed line) and 98% (63 out of 64) exhibited a liver expression value that was at least the median value of all tissue expression for a protein (**Figure 3g**, pie chart insert). These mRNA expression analyses therefore provide further evidence that the hepatocyte secretome is biotinylated by these AAV-*Tbg* viruses.

### Dynamic and nutrient-dependent reprogramming of the hepatocyte secretome

Cellular secretion can be dynamically altered by environmental stressors and other complex physiologic perturbations. Because proximity labeling is initiated by administration of biotin, we reasoned that our strategy could provide sufficient temporal resolution to enable detection of dynamic secretome changes. To test this hypothesis, we analyzed the hepatocyte secretome in response to a two-week period of feeding with high fructose, high sucrose (HFHS) diet^22–24^, a dietary perturbation that powerfully stimulates hepatic lipid accumulation. A cohort of mice was transduced with AAV-*Tbg* virus and, 7 days after transduction, switched to either HFHS diet or maintained on chow diet (**Figure 4a**). Biotin was administered daily from days 14 through 21 (24 mg/kg/day, intraperitoneal injection) and mice were analyzed 24 h after the final injection. Dramatic accumulation of hepatic lipid was verified by Oil Red O staining of liver sections (**Figure 4b**). Next, hepatocyte secretomes were analyzed by anti-biotin blotting of blood plasma. As shown in **Figure 4c-e**, HFHS diet dramatically suppressed biotinylated protein signal in plasma from AAV-*Tbg*-Mem and AAV-*Tbg*-ER constructs by 50-60% from chow-fed mice levels. By contrast, AAV-*Tbg*-Cyto transduced mice exhibited a remarkable 7-fold increase in total plasma biotinylation signal that was largely restricted to the dramatic appearance of a single ∼45 kDa protein (**Figure 4d**). Importantly, we verified that the intracellular biotinylation of hepatocyte proteomes was not statistically different between chow and HFHS mice (**Supplemental Figure 5**), demonstrating that the observed differences in plasma protein biotinylation are due to changes in secretion, rather than intracellular labeling. These data therefore establish that our biotinylation approach can detect dynamic changes to cellular secretomes in response to complex physiologic perturbations.

**Figure 4.**
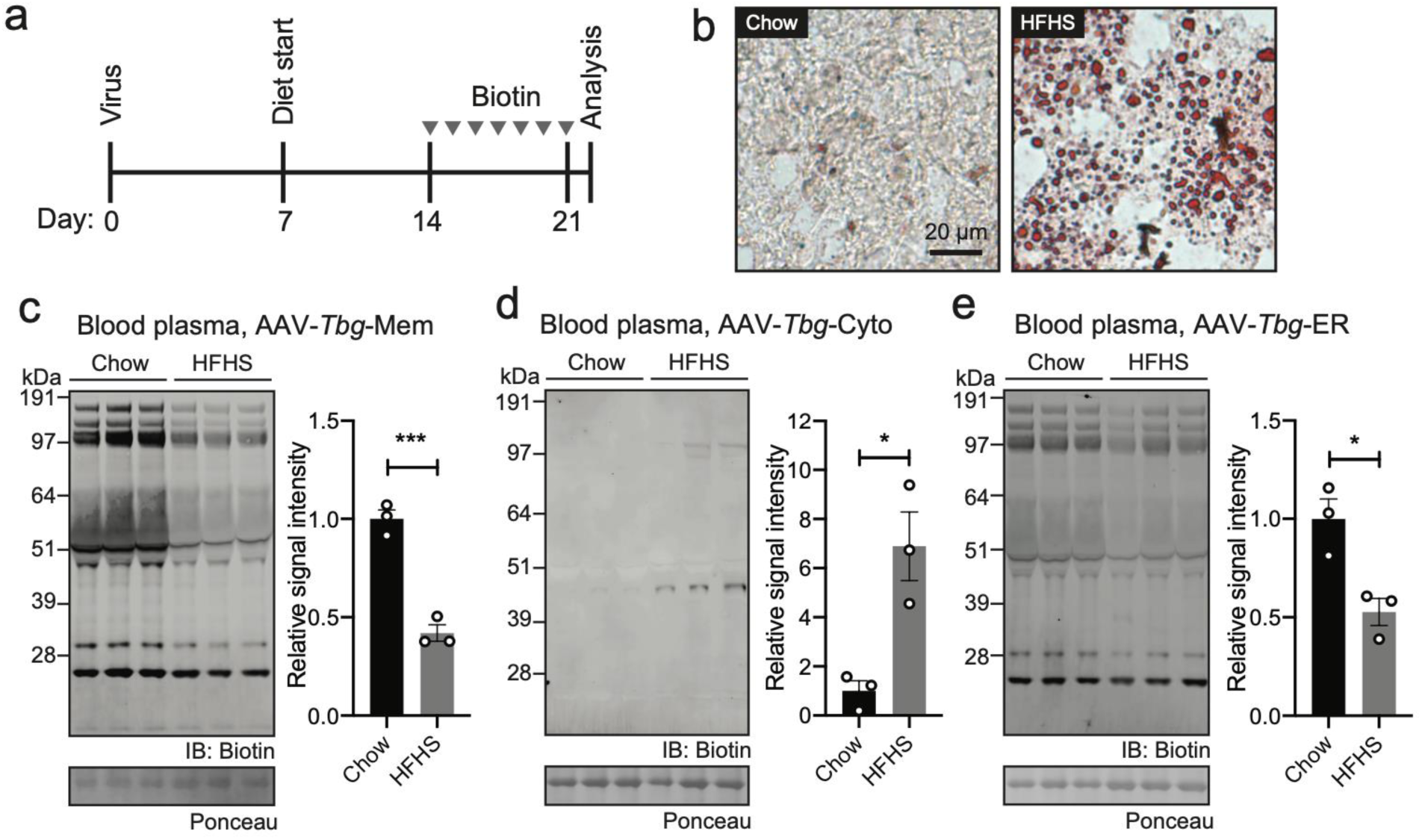
Nutrient-dependent dynamic modulation of the hepatocyte secretome. **(a)** Schematic diagram of the time course of this dietary perturbation experiment. Grey arrows indicate daily biotin administration (24 mg/kg, intraperitoneally). **(b)** Representative Oil Red O staining of fresh frozen, OCT embedded liver sections from mice fed chow or HFHS diet for two weeks. **(c-e)** Anti-biotin (top gel), Ponceau staining (bottom gel), and total band intensity quantitation (right panels) of streptavidin-purified plasma from mice transduced with AAV-*Tbg*-Mem **(c)**, AAV-*Tbg*-Cyto **(d)**, or AAV-*Tbg*-ER **(e)** viruses and fed either chow or HFHS diet for two weeks. Biotin was administered daily for the final week of diet treatment (24 mg/kg/day, intraperitoneally) and blood plasma was harvested 24 h after the final biotin injection. For (**c-e**), data are shown as means ± SEM, N = 3/group. * P < 0.05, *** P < 0.001.

### Unconventional secretion of hepatocyte BHMT

The observation that high fructose, high sucrose feeding stimulates secretion of a single 45 kDa polypeptide as determined by AAV-*Tbg*-Cyto labeling was entirely unexpected. The selectivity for a single polypeptide instead of the entire proteome strongly suggested a highly regulated secretion event. Furthermore, labeling of the 45 kDa polypeptide by AAV-*Tbg*-Cyto and not the other luminally-oriented constructs suggested that export event might be mediated via an unconventional secretory pathway. To determine the identity of this 45 kDa protein, we transduced a new cohort of mice with AAV-*Tbg*-Cyto (N = 3/group). As control groups, we also included a parallel cohort group of mice without virus transduction (“control,” N = 3, for 9 mice total). HFHS diet and biotin administration were performed exactly as described above and biotinylated plasma proteins were purified on streptavidin beads and analyzed by LC-MS/MS (**Supplemental Table S3**). Only a single protein was statistically different by ANOVA and enriched in HFHS versus control mice: the cytosolic enzyme BHMT (betaine-homocysteine S-methyltransferase, **Figure 5a**). BHMT has a predicted molecular weight of 45 kDa, precisely matching the expected molecular weight by gel-based analysis. By quantitation of eluting peptides, plasma BHMT protein was 4-fold elevated in streptavidin-purified plasma from AAV-*Tbg* (HFHS) versus AAV-*Tbg* (chow) mice (**Figure 5b**), a magnitude of change that also largely matched that observed by gel-based quantitation. A representative chromatogram for a tryptic peptide corresponding to BHMT is shown in **Figure 5c**. We therefore conclude that HFHS diet induces a previously undescribed reprogramming of hepatocyte secretion characterized by global suppression of classical secretion pathways with concomitant and selective increase in unconventional BHMT protein secretion.

**Figure 5.**
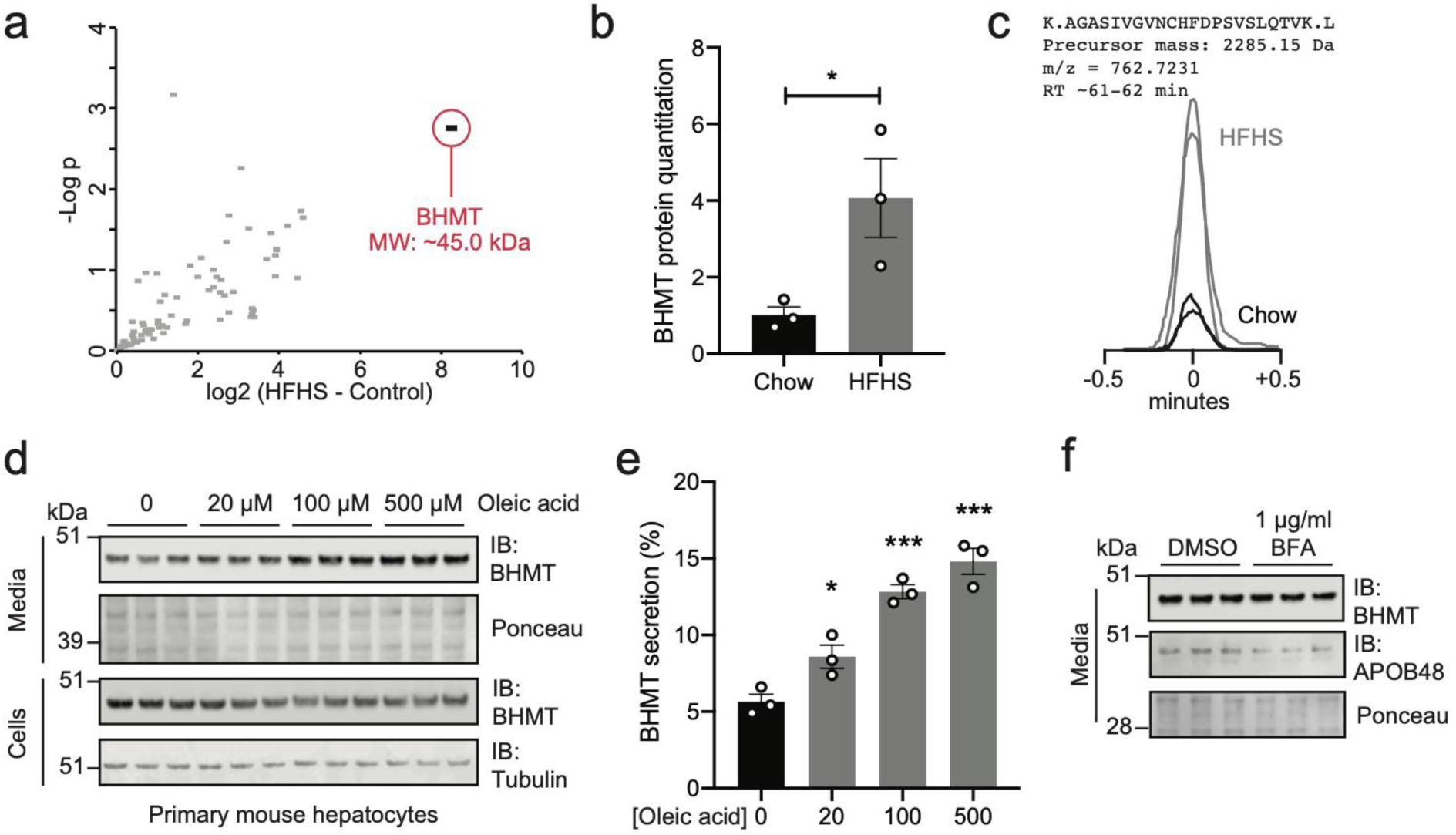
Unconventional secretion of hepatocyte BHMT. **(a)** Proteomic analysis of streptavidin-purified plasma proteins from AAV-*Tbg*-Cyto (HFHS) versus control (chow) mice. BHMT protein is highlighted in red. **(b)** Quantitation of streptavidin-purified plasma BHMT protein fold change levels in AAV-*Tbg*-Cyto (HFHS) and AAV-*Tbg*-Cyto (chow) samples relative to control (chow) samples by LC-MS/MS. **(c)** Extracted ion chromatograms of a representative tryptic peptide from streptavidin-purified plasma BHMT in AAV-*Tbg*-Cyto (HFHS) and AAV-*Tbg*-Cyto (chow). **(d, e)** Anti-BHMT blotting (**d**) and quantification of BHMT protein levels (**e**) in cell lysates or conditioned media from primary mouse hepatocytes isolated from 6 to 12-week old male C57BL/6J mice. Where indicated, hepatocytes were treated with 20 μM, 100 μM or 500 μM oleic acid in serum-free media for 4 h. **(f)** Anti-BHMT and anti-APOB48 blotting in conditioned media of primary mouse hepatocytes isolated from 6 to 12-week old male C57BL/6J mice treated with DMSO or 1 μg/ml brefeldin A in serum-free media for 4h. For **(b)** and **(e),** data are shown as means ± SEM, N = 3/group. * P < 0.05, *** P < 0.001. For (**d-f**), experiments were performed two times and similar results were obtained.

From publicly available single cell murine expression datasets^25^, *Bhmt* mRNA expression is nearly exclusive to the liver (**Supplemental Figure 6**). BHMT has also been previously characterized as a cytosolic enzyme involved in choline metabolism, hepatic lipid accumulation, and hepatocellular carcinogenesis^26^. However, BHMT was not previously known to be secreted. We therefore sought to better characterize BHMT secretion in a cell culture system. Primary mouse hepatocytes were isolated and BHMT protein was measured by Western blotting. As shown in **Figure 5d**, BHMT was robustly detected in both cell lysate and conditioned media even under basal conditions. Addition of increasing concentrations of oleic acid to induce intrahepatocellular lipid accumulation produced a dose-dependent increase in extracellular BHMT with concomitant decreases of intracellular levels (**Figure 5d**). Quantification of band intensities revealed a ∼3-fold stimulation of BHMT secretion at the highest doses of oleic acid in vitro (**Figure 5e**). To confirm that BHMT secretion was not mediated by conventional secretory pathways, primary mouse hepatocytes were treated with brefeldin A (1 µg/ml) to block classical ER/Golgi-mediated trafficking. Under these conditions in the media, extracellular BHMT levels were unchanged whereas levels of the classical secreted protein APOB was reduced (**Figure 5f**). Lastly, transfection of flag-tagged BHMT in HEK293T cells revealed that all transfected BHMT was retained intracellularly even with oleic acid treatment (**Supplemental Figure 7**), demonstrating a surprising cell type specificity for BHMT secretion from hepatocytes, but not HEK293T cells. We therefore conclude that BHMT is secreted from primary hepatocytes via an unconventional secretory pathway.

### A secretome atlas across four diverse cell types in mice

Lastly, we sought to determine whether this labeling approach could be applied to cell types beyond hepatocytes. Three additional cell types (myocytes, pericytes, and myeloid cells) were selected for in vivo secretome analysis. This diverse panel of cell types included cells restricted to one organ (e.g., myocytes) as well as those exhibiting anatomically diverse localizations (pericytes and myeloid). For these studies, the ER-TurboID cassette was used based on our previous observations that this single construct yielded the broadest secretome coverage. To target myocytes, we generated a proximity biotinylation AAV construct driven by the triple myosine creatine kinase promoter^27^ (*tMCK*, **Supplemental Figure 8a,b**). For targeting pericytes and myeloid cells, a conditional cre-dependent FLEx^28^ AAV strategy was employed (**Supplementary Figure 8c,d**). In this strategy, cre-dependent inversion and expression of the proximity biotinylation FLEx cassette in pericytes or myeloid cells was induced by systemic viral transduction and subsequent tamoxifen injection of *Pdgfrb*-creERT2 or *LysM*-creERT2^29, 30^ mice, respectively (see **Methods**). Lastly and as a positive control, the AAV-FLEx viruses were also transduced into *Albumin*-cre mice^31^ to label the hepatocyte secretome (**Supplementary Figure 8e,f**). All three mouse cre driver lines as well as the tMCK promoter had been previously validated to be highly specific for their respective cell types^29–31^. Following viral transduction and labeling by biotin-supplemented water (N=3 animals per cell type), biotinylated proteins isolated from blood plasma were enriched on streptavidin beads and processed for shotgun LC-MS/MS analysis.

In total, 5,109 peptides corresponding to 446 proteins were detected in this experiment (**Supplemental Table 4**). To identify cell type-specific secreted proteins, we filtered the proteomic dataset to identify proteins that were (1) detected across all three biological replicates in at least one cell type and also (2) determined to have statistically different expression (by ANOVA) in at least one condition. 240 proteins passed these filtering criteria. As expected, principal component analysis demonstrated a clear separation of the secretomes based on cell type (**Supplemental Figure 9**). As a positive control, we compared the hepatocyte secretome obtained in this experiment to that obtained using AAV-*Tbg* virus. Despite genetic and technical differences between the two experiments (e.g., using the *Tbg* promoter versus *Albumin*-cre, intraperitoneal biotin injection versus biotin-supplemented water), a majority (74%) of the *Tbg* hepatocyte secretome was recovered in these *Albumin*-cre datasets (**Supplemental Figure 10**), a finding that both validates the conditional cell type-selective strategy used as well as the robustness of this secretome profiling approach.

Next, hierarchical clustering of the Z-score normalized intensities was next used to visualize the data in a manner that enabled visualization of both proteins with enriched secretion by one cell type as well as proteins commonly secreted by multiple cell types (**Figure 6a**). This clustering revealed that different secreted proteins could distinguish between each of the four cell types (**Figure 6a,b**). For instance, myostatin (GDF8), a well-established myokine^32, 33^, was also identified here as a myocyte secretome-selective polypeptide (**Figure 6b**). Similarly, myeloid secretomes were selectively enriched in the coagulation component F13A; pericyte secrettomes were enriched in diverse immunoglobulins (e.g., KV5A6), and hepatocytes selectively secreted apolipoproteins (e.g., APOH) (**Figure 6b**). This comparative analysis also identified secreted proteins common to multiple cell type secretomes, such as VWF (selectively absent in hepatocyte secretomes), members of the serine protease inhibitor A1 family (AT1AT members, selectively absent in the myeloid cell secretome), and TGM2 (selectively absent in the myocyte secretome) (**Figure 6c**). Lastly, our secretome atlas provides direct in vivo evidence for cellular sources of plasma proteins that are not classically associated with those cell types. For instance, the adipocyte hormone adiponectin^34, 35^ (ADIPOQ) clustered with myostatin and was selectively enriched in the myocyte secretome versus the other cell types (**Figure 6d**). We interpret this observation as biochemical evidence for muscle as a non-adipose origin for circulating adiponectin, a finding supported by previous studies identifying adiponectin secretion from both myocytes and cardiomyocytes in culture^36, 37^. Taken together, these data establish the in vivo secretomes of four different cell types directly from blood plasma of mice, identify secreted proteins that distinguish between cell types in vivo, and uncover unexpected cellular sources for classical plasma proteins.

**Figure 6.**
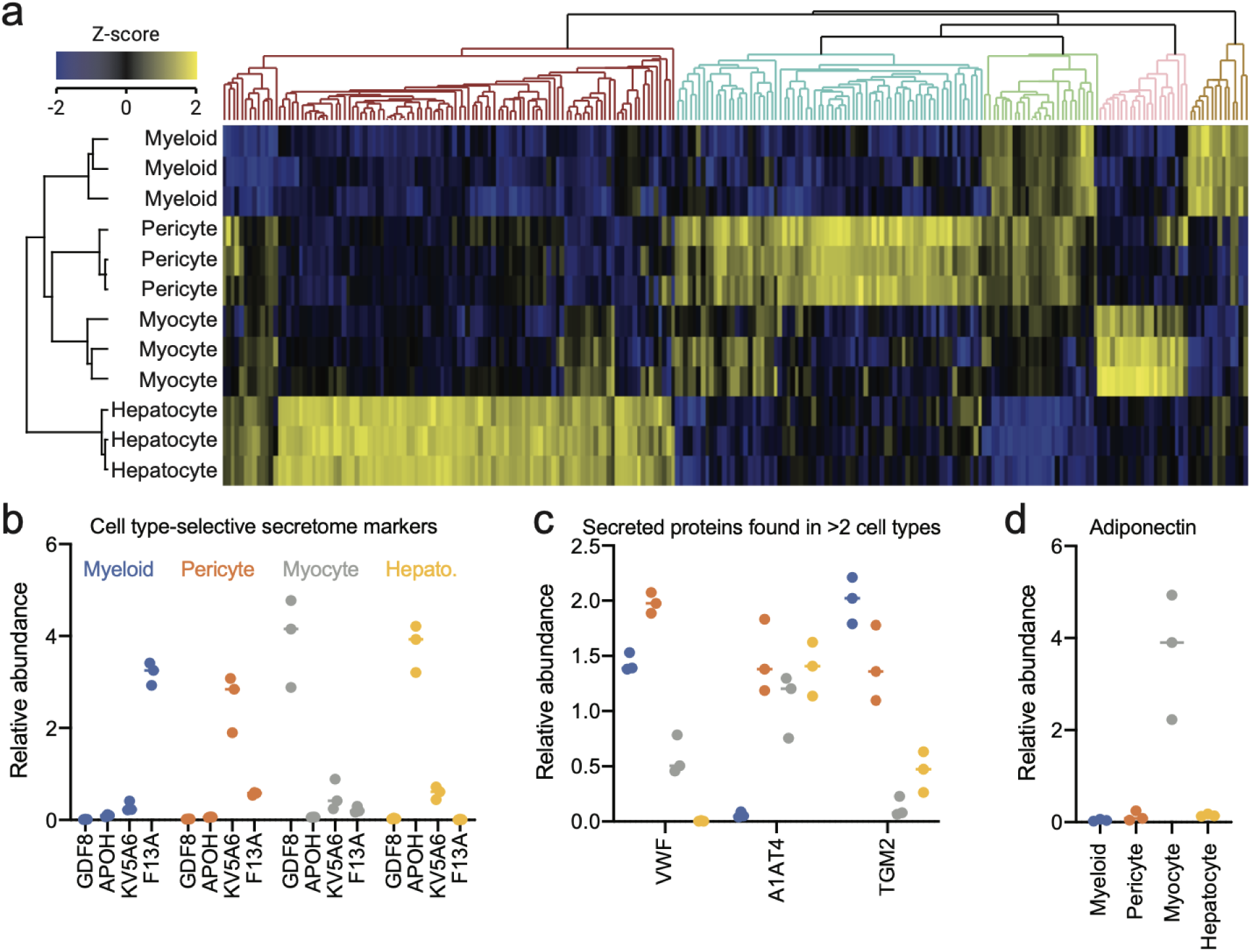
A secretome atlas of four cell types in vivo. **(a)** Hierarchical clustering of Z-score intensities across all differentially detected streptavidin-purified plasma proteins from myeloid, pericyte, myocyte, and hepatocyte secretomes in vivo. (**b-d**) Relative protein abundance of the indicated secreted protein in each of the four secretomes. For (**a-d**), the following AAVs and mouse lines were used: myeloid (*LysM*-creERT2, AAV-FLEx); pericyte (*Pdgfrb*-creERT2, AAV-FLEx), myocyte (wild type, AAV-*tMCK*); hepatocyte (*Albumin*-cre, AAV-FLEx). For viral transduction of myocyte secretomes, wild-type C57BL/6J pups (age P2) were injected with AAV-tMCK virus (10e11 GC/mouse, intraperitoneally) and biotin labeling was initiated at 6 weeks of age. For transduction of hepatocyte secretomes, AAV-FLEx virus (10e11 GC/mouse, intravenously) was injected into hemizygous 6-8 week old male *Albumin*-cre mice and biotin labeling was initiated after a one week transduction period. For transduction of myeloid and pericyte secretomes, AAV-FLEx virus (10e11 GC/mouse, intravenously) was injected into 6 to 8- week old hemizygous *Pdgfrb*-creERT2 or *LysM*-creERT2 male mice. After a one week transduction period, tamoxifen was administered daily for five days (2 mg/mouse/day, intraperitoneal) and biotin labeling was initiated one week after the final tamoxifen injection. For all groups, biotin was administered via biotin-supplemented water (0.5 mg/ml, 3 days). N = 3 mice/group.

## DISCUSSION

Here we introduce a bio-orthogonal strategy for direct detection, enrichment, and profiling of cell type-selective secretomes in mice. This approach relies on proximity biotinylation of intracellular proteins and following those proteins as they are secreted into blood plasma. By using a suite of cell type-specific viral biotinylation reagents, we generate a proteomic atlas of four cell type secretomes by directly purifying biotinylated proteins from blood plasma in mice. Our strategy represents the first direct biochemical detection of cell type-selective secretomes in an intact mammalian organism. Furthermore, our in vivo secretome atlas validated known cell type-protein pairs (e.g., GDF8 secreted by myocytes; FETUB from hepatocytes), uncovered sets of secreted proteins that could distinguish between cell types, and identified new cellular origins for classical plasma proteins (e.g., hepatocytes as source of shed LIFR and EGFR ectodomains; myocytes as a source of adiponectin).

Importantly, our in vivo profiling approach also enables mapping of dynamic secretome changes to complex physiologic perturbations. By profiling the in vivo hepatocyte secretome after high fructose high sucrose feeding, we uncover a dynamic modulation of hepatocyte secretion characterized by selective stimulation of unconventional secretion and broad suppression of classical secretion pathways. Previously, HFHS diets has been reported to either increase^38–40^ or suppress^41^ the secretion of hepatocyte-derived plasma proteins, with individual polypeptides typically used as examples. The in vivo hepatocyte secretome data here further contextualizes these past individual observations by providing direct and unbiased biochemical evidence that hepatic lipid accumulation bidirectionally alter distinct secretory pathways, depending on the secretory substrate and pathway of export.

An entirely unexpected observation from our secretome profiling was the identification of the cytosolic enzyme BHMT as a lipid-induced unconventionally secreted protein from hepatocytes. BHMT secretion can be recapitulated in primary murine hepatocytes but not in other cells (e.g., transfected HEK293T cells), providing evidence for a cell-type specific regulation of its secretion. At this time, the precise pathway for BHMT secretion remains unknown, though selective labeling by AAV-*Tbg*-Cyto in vivo and inhibitor experiments in vitro suggest that this secretion pathway is not mediated by the ER/Golgi. Importantly, while the intracellular functions of BHMT have been well-established, its extracellular roles and potential as a biomarker for fatty liver disease remain critical questions for future studies.

Lastly, we estimate that this secretome profiling approach exhibits a sensitivity of ∼100 pg/ml based on the identification of several very low abundance hormones (e.g., GDF8 “myostatin”, ∼10 ng/ml) and ectodomain shed receptors (e.g., LIFR, ∼100 pg/ml) **(Supplementary Figure 11**). This sensitivity is similar to that of other state-of-the-art technologies currently available for multiplexed plasma proteomics (e.g., aptamer-based SOMAscan^42, 43^, ∼ng/ml) and is a dramatic ∼3-log improvement compared to standard shotgun plasma proteomics approaches (sensitivity at ∼µg/ml). We attribute this high level of sensitivity to be due to the enrichment provided by local biotinylation of secreted proteins. In the future, additional improvements in sensitivity to reach that of individual immunoassays (∼pg/ml) could potentially be achieved by the development of more efficient in vivo proximity biotinylation reagents and/or the use of targeted proteomic pipelines rather than the data-dependent acquisition approaches used here.

Projecting forward, an important next step will be to expand this secretome technology to additional cell types using either adeno-associated viruses or conditional mice. By providing global biochemical portraits of cell type-specific secretomes and their contribution to the circulating polypeptides in plasma, these studies pave the way for cell type-specific dissection of the plasma proteome and discovery of additional polypeptide mediators of intercellular communication and organismal homeostasis.

## METHODS

### Cell lines cultures

HEK 293T cells were obtained from ATCC (CRL-3216) and cultured in complete media consisting of DMEM with L-glutamine, 4.5 g/l glucose and sodium pyruvate (Corning 10013CV) supplemented with 10% FBS (Corning 35010CV), and 1:1000 penicillin/streptomycin (Gibco 15140-122). Cell were at 37 °C, 5% CO2 for growth. For transient transfection, cells were transfected in 10 cm^2^ or 6-well plates at ∼60% confluency using PolyFect (Qiagen 301107) and washed with complete media 6 h later.

### General animal information

Animal experiments were performed according to procedures approved by the Stanford University IACUC. Mice were maintained in 12 h light-dark cycles at 22°C and fed a standard irradiated rodent chow diet. Where indicated, high sucrose high fructose diet (Research Diets D09100310) was used. C57BL/6J male mice (stock number 000664), homozygous *Albumin*-cre male mice (stock number 003574), homozygous *Pdgfrb*-P2A-CreERT2 male mice (stock number 030201), hemizygous *LysM*-CreERT2 male mice (stock number 031674), and FVB/NJ female mice (stock number 001800) were purchased from Jackson Laboratories. *Albumin*-cre and *Pdgfrb*-P2A-creERT2 male mice were crossed with female C57BL/6J mice to generate hemizygous *Albumin*-cre and *Pdgfrb*-P2A-creERT2 mice respectively. *LysM*-creERT2 male mice were crossed with FVB/NJ female mice to generate hemizygous *LysM*-creERT2 mice. Genotypes were verified following genotyping protocols and using primers listed on Jackson Laboratories website.

### Materials

The following antibodies were used: anti-V5 antibody (Invitrogen R960-25), mouse anti-FLAG M2 antibody (Sigma F1804), anti-beta Actin (Abcam ab8227), anti-beta Tubulin antibody (Abcam ab6046), anti-Albumin (Abcam ab19194), anti-BHMT (Abcam 96415), anti-mouse IgG IRDye 680RD (LI-COR 925-68070), anti-rabbit IgG IRDye 800RD (LI-COR 926-32211), anti-biotin Streptavidin AlexaFluor-680 (ThermoFisher S32358), anti-mouse IgG AlexaFluor-488 (Life Technologies A11029-EA. The following plasmids were used: mouse PM20D1-flag (Addgene 84566). pENN.AAV.tMCK.PI.eGFP.WPRE.bGH (Addgene 105556), AAV.TBG.PI.Cre.rBG (Addgene 107787-AAV8), pCAG-Cre (Addgene 13775), V5-TurboID-NES_pCDNA3 (Addgene 107169), human FGF1-myc-DDK (Origene RC207434), human flag-FGF2 (SinoBiological HG10014-NF). ER-TurboID and Mem-TurboID were synthesized (IDT) and inserted into pcDNA3, pENN.AAV.tMCK.PI.eGFP.WPRE.bGH, and AAV.TBG.PI.Cre.rBG expression plasmids with pENTR-d-topo cloning (ThermoFisher K240020) and restriction site cloning. mBHMT-flag was synthesized (IDT) and inserted into pcDNA-DEST40 expression plasmids with pENTR-d-topo cloning. The following adeno-associated viruses (and batch numbers in parentheses) were produced at the Penn Vector Core: AAV8-*Tbg*-Mem (V7155S), AAV8-*Tbg*-Cyto (V7119S), AAV8-*Tbg*-ER (V7069S), AAV9-*tMCK*-ER (V7074S), AAV8-CB7-FLEx (V7258S).

### Secretome labeling in cell culture

24 h after transfection, HEK293T cells were washed twice with PBS and incubated with serum-free media containing 500 μM biotin. 18 h later, labeling was terminated by washing cells five times with ice-cold PBS. Conditioned media was collected and concentrated 20-fold using 10 kDa filter tubes (Millipore UFC801024). Cells were harvested and lysed by probe sonication in RIPA buffer consisting of 1% NP40, 0.1% SDS, 0.5% sodium deoxycholate and 1:100 HALT protease inhibitor (ThermoFisher 78429). Cell lysates were centrifuged at 13,000 rpm for 10 min at 4°C. The supernatant was isolated, boiled for 10 min at 95°C in 4x NuPAGE LDS Sample Buffer supplemented with 100 mM DTT, and analyzed by Western blot. For validation of AAV8-FLEx constructs in HEK293T cells, cells were maintained in culture for 3 days after co-transfection of pCAG-Cre and AAV8-FLEx plasmids with media refreshed every day to allow sufficient recombination before biotin addition. Labeling was initiated as described above.

### Determination of biotinylation efficiency in cell culture

HEK293T cells were transfected, labelled and harvested as described above. 2 ml conditioned media was collected and concentrated 10x using 10 kDa filter tubes (Millipore UFC801024) to 200 µl. 10 µl conditioned media was saved as input fraction. 100 μl Dynabeads MyOne Streptavidin T1 magnetic beads (ThermoFisher 65602) were washed twice with washing buffer (50 mM Tris-HCl, 150 mM NaCl, 0.1% SDS, 0.5% sodium deoxycholate, 1% NP40, and 1 mM EDTA, 1x HALT protease inhibitor, 5 mM Trolox, 10 mM sodium azide, and 10 mM sodium ascorbate), resuspended in 190 μl conditioned media and incubated at 4°C overnight with rotation. 200 μl supernatant was collected as flow through fraction. The beads were subsequently washed thoroughly twice with 1 ml of RIPA lysis buffer and eluted by boiling at 95C for 10min in 20 μl of 2x sample buffer supplemented with 20 mM DTT and 2 mM biotin. 25 µl eluate was finally collected. 5% of input, elution and flow through were analyzed by Western blotting using anti-flag and anti-biotin antibodies. Since PM20D1-flag and FGF1-flag were used for transfection, eluate was considered as both anti-flag and anti-biotin positive fraction. The band intensity was calculated with ImageJ. The total biotinylated flag-tagged protein is calculated as total(biotinylated) = anti-flag(elute)*[anti-biotin(flow through) + anti-biotin(elute)]/anti-biotin(elute). The biotinylation efficiency is calculated as total(biotinylated)/[anti-flag(flow through) + total(biotinylated)].

### Western blot analysis

For all the gels shown in this study, proteins separated on NuPAGE 4-12% Bis-Tris gels were transferred to nitrocellulose membrane. Media and plasma samples were further incubated with Ponceau S solution to ensure equal loading and washed with PBST (PBS, 0.05% Tween 20). Blots were blocked in Odyssey blocking buffer for 30 min at room temperature and incubated with primary antibodies (Mouse anti-V5 antibody, Invitrogen R960-25, 1:1000 dilution; Mouse anti-Flag antibody (Sigma F1804), 1:1000 dilution; Rabbit anti-beta Tubulin antibody (Abcam ab6046), 1:5000 dilution; Rabbit anti-beta-Actin (Abcam ab8227), 1:5000 dilution; Rabbit anti-BHMT antibody (Abcam 96415); in blocking buffer for 1 h at room temperature or overnight at 4°C. After washing three times with PBST, blots were stained with species matched secondary antibodies (Streptavidin-AlexaFluor680, ThermoFisher S32358, 1:1000 dilution; Goat anti-mouse IRDye 680RD, LI-COR 925-68070, 1:10000 dilution; Goat anti-rabbit IRDye 800RD, LI-COR 925-68070, 1:10000 dilution. Blots were then washed three times with PBST and imaged with Odyssey CLx Imaging System.

### Immunofluorescence of transfected cells

Glass coverslips were immerged into 1M HCl on a shaker overnight, washed twice with distilled water, once with EtOH and air dried. Glass coverslips were then incubated with 5 µg/ml fibronectin (Millipore 341635-1MG) in 6-well plate at 37 °C for 1 h. Fibronectin solution was then removed and plates were washed twice with distilled water and once with complete media. Transfected HEK293T cells were trypsinized and replated on coated coverslips in 6-well plate and incubated in complete media overnight. Cells were washed twice with PBS and fixed in 4% formaldehyde in PBS (w/v) (ThermoFisher PF18620910) for 15 min at room temperature. Cells were then washed twice with PBS and permeabilized with methanol: acetone (1:1) for 2 min at room temperature. Cells were again washed twice with PBS and incubated with Odyssey blocking buffer for 1 h at room temperature. Blocked cells were then stained with primary antibodies (Goat anti-mouse AlexaFluor488, Life Technologies A11029-EA, 1:1000 dilution) in blocking buffer for 30 min and with 1 μg/ml DAPI in blocking buffer for 10 min. Cells were washed twice with PBS, mounted and imaged with confocal fluorescence microscopy. Images were taken with a Leica TCS SP5 AOBS confocal microscope equipped with a laser scanner and a 63x oil immersion objective.

### Viral transduction

For AAV-*Tbg*-Mem and AAV-*Tbg*-Cyto, 6 to 8-week old male mice (C57BL/6J) were injected via tail vein with 29 G syringe (ThermoFisher 14-841-32) at a dose of 10e11 GC/mouse diluted in saline in a total volume of 100 μl/mouse. AAV-*Tbg*-ER was performed identically but at a titer of 10e10 GC/mouse. For AAV-*tMCK* virus injections, post-natal day 2 pups (C57BL/6J) were injected Intraperitoneally with 31 G syringe (BD 328290) at a dose of 10e11 GC/mouse diluted in saline in a total volume of 50 μl/mouse. One week after transduction of AAV-*Tbg* viruses or eight weeks after transduction of AAV-*tMCK* viruses, biotin labeling was initiated as described below.

For transduction of conditional cre mice, AAV-FLEx virus (10e11 GC/mouse) was injected into 6 to 8-week old hemizygous *Pdgfrb*-P2A-creERT2 or *LysM*-creERT2 male mice. After a one-week transduction period, tamoxifen (Sigma, T5648-1G) was prepared as a 20 mg/ml solution in corn oil and administered daily for five days (100 μl/day, intraperitoneal) to induce recombination. After the final tamoxifen injection, mice were housed in their home cages for one additional week before biotin labeling experiments.

### Biotin labeling in mice

Biotin was administered by injection (24 mg/ml, intraperitoneally, in a solution of 18:1:1 saline:kolliphor EL:DMSO, final volume 200 µl/mouse/day) for three consecutive days. For labeling with biotin in drinking water, the water was supplemented with biotin (0.5 mg/ml) for three consecutive days.

### Plasma and tissue sample preparation from mice

24 h after the final biotin dose, blood was collected using 21G needle (BD 305129) by submandibular bleed into lithium heparin tubes (BD 365985) and immediately spun (5,000 rpm, 5 min, 4°C) to isolate the plasma fractions. All tissues were dissected, collected into Eppendorf tubes, and immediately frozen on dry ice and stored in −80°C. Tissues were homogenized in 0.5 ml of cold RIPA buffer using a Benchmark BeadBlaster Homogenizer at 4°C. The mixture was centrifuged (13,000 rpm, 10 min, 4°C) to spin out beads and the homogenate was quantified and analyzed by Western blot. To remove excess biotin in samples, 200 μl plasma from a single animal was diluted in 15 ml PBS and concentrated 30-fold using 3 kDa filter tubes (Millipore UFC900324) at 4,000 rpm for 1.5 h. The flow-through was removed and the concentration step was repeated until a 500 μl final solution was recoved at 9000-fold final dilution. To enrich biotinylated materials from proteomic samples, 200 μl Dynabeads MyOne Streptavidin T1 magnetic beads (ThermoFisher 65602) were washed twice with washing buffer (50 mM Tris-HCl, 150 mM NaCl, 0.1% SDS, 0.5% sodium deoxycholate, 1% NP40, and 1 mM EDTA, 1x HALT protease inhibitor, 5 mM Trolox, 10 mM sodium azide, and 10 mM sodium ascorbate) and resuspended in 100 μl washing buffer. Plasma solution was added to beads and incubated at 4°C overnight with rotation. The beads were subsequently washed thoroughly twice with 1 ml of RIPA lysis buffer, once with 1 ml of 1 M KCl, once with 1 ml of 0.1 M Na2CO3, once with 1 ml of 2 M urea in 10 mM Tris-HCl (pH 8.0), and twice with 1 ml washing buffer. Biotinylated proteins were then eluted from the beads by boiling the beads at 95C for 10min in 60 μl of 2x sample buffer supplemented with 20 mM DTT and 2 mM biotin. Successful enrichment was validated by running 5 ul sample elution on NuPAGE 4-12% Bis-Tris gels followed by silver stain (ThermoFisher LC6070) following the manufacturer’s protocol.

### Proteomic sample processing

60 μl of eluted and boiled streptavidin-purified plasma samples were cooled to room temperature for 3 min and digested using a mini S-trap protocol provided by the manufacturer (Protifi), described as follows^44^. Cysteines were alkylated using iodoacetamide (Sigma A3221) added to a final concentration of 30 mM and incubated in the dark at room temperature for 30 min. Samples were acidified with phosphoric acid (to 1.2% final volume, vortexed to mix) and 420 μl 100 mM TEAB in 90% MeOH was added to each. Samples were loaded onto micro S-trap columns, flow through was discarded, and the centrifugation step was repeated until all the solution passed through the column. Following washing with 100 mM TEAB in 90% methanol, 1 μg trypsin (Promega) was added to the S-trap for 90 minutes at 47 °C. After trypsinization, peptides were eluted from S-traps with 50 mM TEAB (40 μl), 0.2% formic acid (40 μl), 50% acetonitrile + 0.2% formic acid (40 μl), and a last wash of 0.2% formic acid in water (40 μl) by centrifugation at 1,000 x g for 60 s. Eluted peptide samples were lyophilized, resuspend in 0.2% formic acid and analyzed by LC-MS/MS.

### LC-MS/MS

All samples were resuspended in 20 μl 0.2% formic acid in water and 5 μl were injected on column for each sample. Peptides were separated over a 50 cm EasySpray reversed phase LC column (75 µm inner diameter packed with 2 μm, 100 Å, PepMap C18 particles, ThermoFisher). The mobile phases (A: water with 0.2% formic acid and B: acetonitrile with 0.2% formic acid) were driven and controlled by a Dionex Ultimate 3000 RPLC nano system (ThermoFisher). An integrated loading pump was used to load peptides onto a trap column (Acclaim PepMap 100 C18, 5 um particles, 20 mm length, ThermoFisher) at 5 µL/min, which was put in line with the analytical column 6 minutes into the gradient for the total protein samples. Gradient elution was performed at 300 nL/min. The gradient increased from 0% to 5% B over the first 6 minutes of the analysis, followed by an increase from 5% to 25% B from 6 to 86 minutes, an increase from 25% to 90% B from 86 to 94 minutes, isocratic flow at 90% B from 94 to 102 minutes, and a re-equilibration at 0% for 18 minutes for a total analysis time of 120 minutes. Precursors were ionized using an EASY-Spray ionization source (ThermoFisher) source held at +2.2 kV compared to ground, and the column was held at 45 °C. The inlet capillary temperature was held at 275 °C. Survey scans of peptide precursors were collected in the Orbitrap from 350-1500 Th with an AGC target of 1,000,000, a maximum injection time of 50 ms, and a resolution of 120,000 at 200 m/z. Monoisotopic precursor selection was enabled for peptide isotopic distributions, precursors of z = 2-5 were selected for data-dependent MS/MS scans for 2 seconds of cycle time, and dynamic exclusion was set to 45 seconds with a ±10 ppm window set around the precursor monoisotope. An isolation window of 0.7 Th was used to select precursor ions with the quadrupole. MS/MS scans were collected using HCD at 30 normalized collision energy (nce) with an AGC target of 50,000 and a maximum injection time of 54 ms. Mass analysis was performed in the Orbitrap with a resolution of 30,000 at 200 m/z and an automatically determined mass range.

### Proteomics data analysis

Raw data were processed using MaxQuant^45^ version 1.6.10.43 and tandem mass spectra were searched with the Andromeda search algorithm^46^. Oxidation of methionine, biotinylation of lysine, and protein N-terminal acetylation were specified as variable modifications, while carbamidomethylation of cysteine was set as a fixed modification. 20 ppm, 4.5 ppm, and 20 ppm were used for first search MS1 tolerance, main search MS1 tolerance, and MS2 product ion tolerance, respectively. Cleavage specificity set to Trypsin/P with 2 missed cleavage allowed. Peptide spectral matches (PSMs) were made against a target-decoy mouse reference proteome database downloaded from Uniprot (17,030 entries). Peptides were filtered to a 1% false discovery rate (FDR), two peptides were required for a protein identification, and a 1% protein FDR was applied. Proteins were quantified and normalized using MaxLFQ^47^ with a label-free quantification (LFQ) minimum ratio count of 2. The match between runs feature was enabled. Quantitative comparisons were performed using Perseus v 1.6.2.2^48^. For quantitative comparisons, protein intensity values were log2 transformed prior to further analysis, proteins with at least two measured values in a condition were retained, and missing values were imputed from a normal distribution with width 0.3 and downshift value of 1.8 (i.e., default values). Significance calculations for pairwise comparisons (**Figure 3**) were performed using a two-tailed t-test with a permutation-based FDR with 250 randomizations, an FDR of 0.05, and an S0 value of 1. For hierarchical clustering, significance was first determined using ANOVA with a permutation-based FDR with 250 randomizations, an FDR of 0.05, and an S0 value of 1. Only significant hits were retained and were then normalized by Z-score calculations before clustering using Euclidean distance. Gene Ontology (GO) term enrichment for **Figure 3** was performed using DAVID^49^ with a background of 500 mouse plasma proteins^50^. For calculating tissue expression data from BioGPS (**Figure 3**), expression profiles across 191 tissues/cells lines that are available in BioGPS were collected for 64 of the 66 enriched proteins in AAV-TBG-Mem and AAV-TBG-ER (two were not available in BioGPS). For each protein, the median expression value across all tissues/cell lines was calculated, as was the total signal observed for a protein (i.e., the sum of signal in all 191 tissues/cells). To generate **Figure 3f**, the fraction of signal for each of the tissues/cell types for a protein was calculated by dividing the tissue/cell type signal by the total signal observed for that protein. Thus, fractions show proportions between 0 and 1, with numbers closer to 1 indicating more signal in a given tissue/cell type. For **Figure 3g**, the signal in liver for each protein was compared to that protein’s media expression value across all tissue/cell types. If signal in liver was greater than or equal to the media expression value, that protein was categorized as having a liver annotation.

### Oil Red O staining of liver sections

Fresh liver tissue was snap frozen as an O.C.T. embedded block (ThermoFisher 23-730-571) and cryosectioned. Frozen slides were fixed with 4% paraformaldehyde for 1 hour. Slides were subsequently washed with 4 changes of ultrapure water, incubated with 60% isopropanol for 5 minutes, incubated with Oil Red O working solution (3:2 Oil Red O: ultrapure water) for 20 minutes (Sigma O1391), and washed with 5 changes of ultrapure water.

### Plasma ALT measurement

20 μl mouse plasma from AAV-*Tbg* viruses transduced mice were used to measure enzymatic activity of circulating alanine transaminase following manufacturer’s protocol (Cayman NC0819930).

### Immunohistochemistry of liver sections

PBS-perfused livers were isolated and post-fixed in 4% (w/v) paraformaldehyde in PBS overnight at 4°C before preservation in 30% (w/v) sucrose in PBS. Livers were sectioned at a thickness of 50 μm on a freezing–sliding microtome, and sections were stored in cryoprotective medium at −20°C. Free-floating sections were blocked with 5% donkey serum for 1.5 hours before overnight incubation at 4°C with primary antibodies (goat anti-Albumin antibody, Abcam ab19194, 1:100 dilution; rabbit anti-V5 antibody, Cell Signaling Technology, 1:500 dilution). Sections were washed, stained with Alexa Fluor-conjugated secondary antibodies (Thermo Fisher Scientific, 1:250 dilution) in blocking buffer for 2.5 hours at room temperature, mounted, and coverslipped with ProLong Gold (Life Technologies) before imaging on a confocal laser-scanning microscope (Zeiss LSM880).

### Isolation and culture of primary mouse hepatocytes

6 to 12 week-old male mice (C57BL/6J) were perfused with perfusion buffer (0.4 g/L potassium chloride, 1 g/L glucose, 2.1 g/L sodium bicarbonate, 0.2 g/L EDTA in HBSS buffer) via cannulate vena cava for 8 min, with digestion buffer (1 mg/ml collagenase IV (Sigma C5138-1G) in DMEM/F12 media) for 8 min. Liver was then removed and passed through 70 μm cell strainer (BD 352350) to obtain crude hepatocytes. Cells were then centrifuged at 50 g for 3 min, resuspended in 10 ml plating media (10% FBS, 2 mM sodium pyruvate, 1% penicillin/streptomycin, 1 μM dexamethasone (Sigma D4902-100MG), 0.1 μM insulin (Sigma I5500) in William E media (Quality Biological 10128-636)) and centrifuged at 50 g for 3 min. The cell pellet was resuspended in 10 ml 45% percoll solution and centrifuged at 100 g for 10 min. This final hepatocyte pellet was then resuspended in 10 ml plating media, centrifuged at 50 g for 5 min and resuspended in 1 ml plating media. Hepatocytes were counted and plated in fibronectin-coated 6-well plate at 1 million cells per well. 4 h later, media were changed into maintenance media (0.2% BSA (Sigma A7906-500G), 2 mM sodium pyruvate, 1% penicillin/streptomycin, 0.1 μM dexamethasone, 1 nM insulin) and incubated overnight with indicated concentrations of oleic acids or DMSO.

### Treatment of cells with oleic acid or brefeldin A

24 h following isolation, primary mouse hepatocytes were washed twice with warm PBS. After aspiration of the second wash, cells were incubated with 2 ml Williams E media containing the indicated concentrations of oleic acid (Sigma 05508-5ML), 1 μg/ml Brefeldin A (Sigma B6542-5MG) or DMSO. 4 h later, cells and conditioned media were collected and processed as previously described. For HEK293T cells, 24 h after transfection, cells were washed with PBS and incubated with serum-free media containing 500 μM oleic acid. 18 h later, cells and conditioned media were collected and processed as previously described.

### Statistics

All measurements were taken from distinct samples. Statistical significance was determined by Student’s two-sided t-test unless otherwise indicated.

## ACKNOWLEDGEMENTS

We thank members of the Long, Bertozzi, Svensson, and Abu-Remaileh labs for helpful discussions. We gratefully acknowledge the staff at the Penn Vector Core for production of adeno-associated viruses. This work was supported by the US National Institutes of Health (DK105203 and DK124265 to JZL and K00CA21245403 to NMR), by the Stanford ChEM-H Institute (DF and CRF), and by the Stanford Diabetes Research Center (P30DK116074).

## AUTHOR CONTRIBUTIONS

WW: Conceptualization, Methodology, Investigation, Writing – Original Draft, Writing – Review & Editing, Visualization

NMR: Methodology, Software, Formal analysis, Investigation, Resources, Data Curation, Writing – Original Draft, Writing – Review & Editing, Visualization

ACY: Investigation

JTK: Investigation

SMT: Investigation

VLL: Investigation

MGC: Investigation

CRB: Methodology, Supervision, Funding acquisition

JZL: Conceptualization, Resources, Writing – Original Draft, Writing – Review & Editing, Supervision, Funding acquisition

## DATA AVAILABILTY STATEMENT

The authors declare that data supporting the findings of this study are available within the paper and its supplementary information files. Figures 3-6 have associated raw data provided in Tables S1-4.

---

**Supplemental Figure 1.**
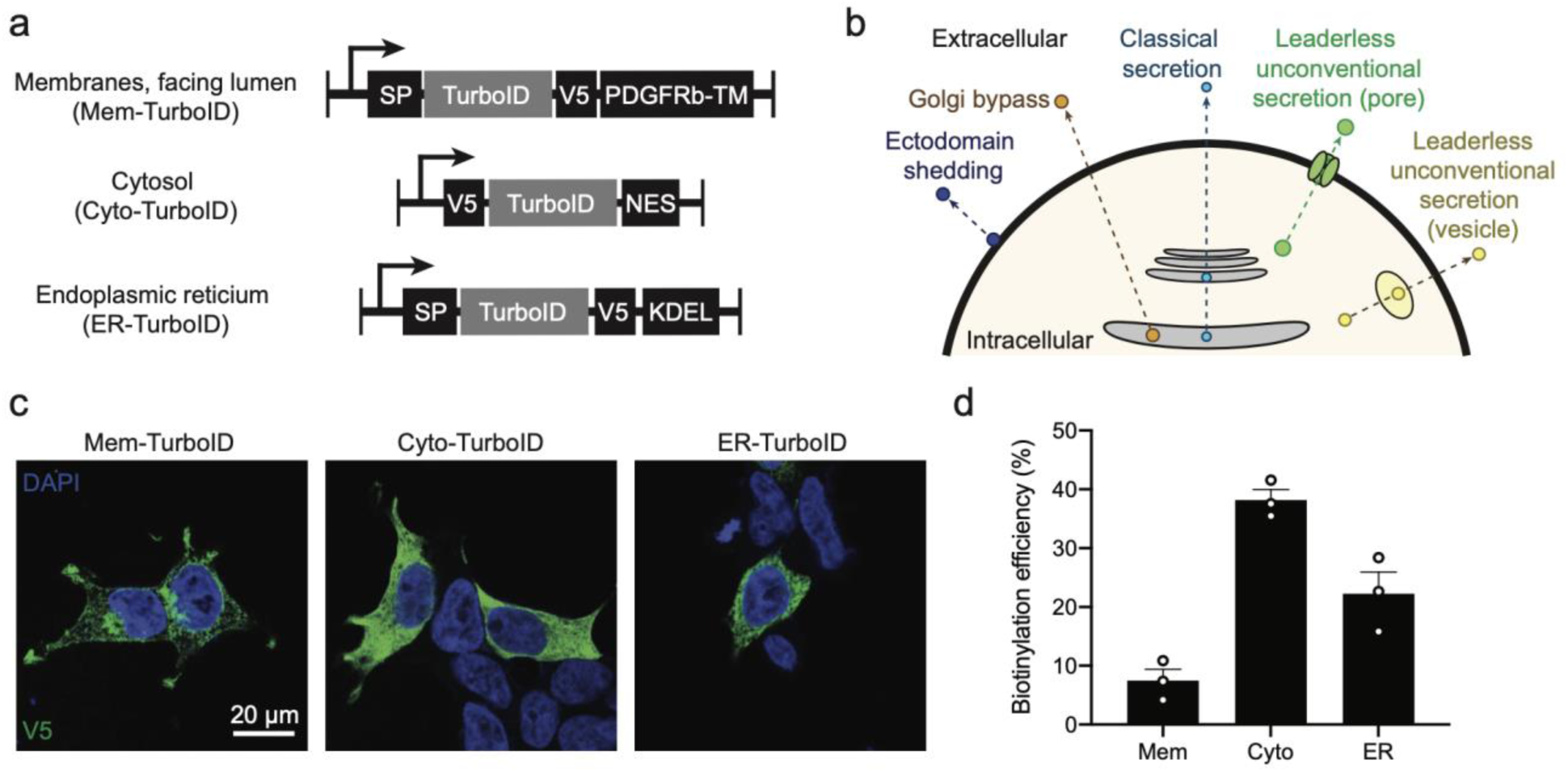
In vitro characterization of proximity labeling constructs. **(a)** Schematic of the proximity labeling constructs used in this study. Abbreviations: SP, signal peptide; PDGFRb-TM, transmembrane domain of PDGFRB receptor; NES, nuclear export signal; KDEL, tetrapeptide ER retention signal. **(b)** Schematic of various pathways for cellular secretion. **(c)** Immunofluorescence using an anti-V5 antibody (green) or nuclei stain (DAPI, blue) of HEK293T cells transfected with the indicated constructs. **(d)** In vitro biotinylation efficiency of Mem-TurboID, Cyto-TurboID, or ER-TurboID by measurement of biotinylated PM20D1-flag or biotinylated FGF1-flag versus the total flag signal in conditioned media. Data are shown as means ± SEM, N = 3/group.

**Supplemental Figure 2.**
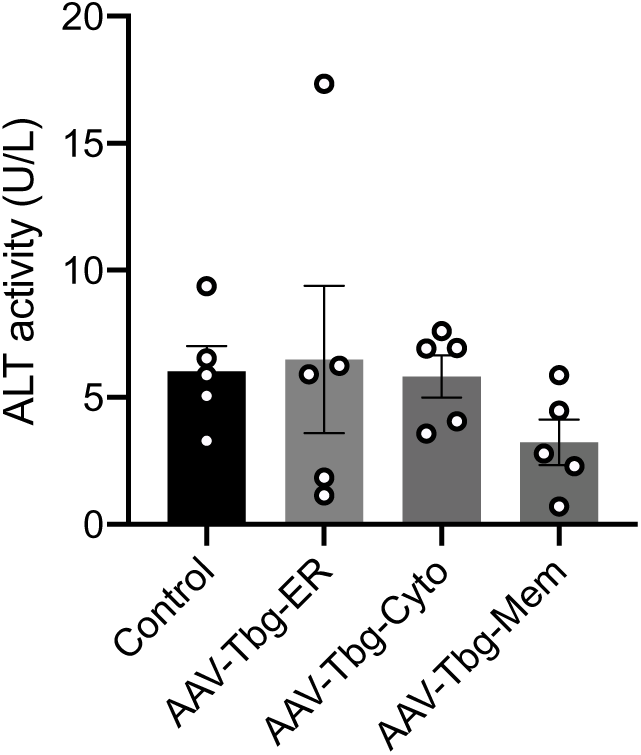
ALT levels in mice transduced with AAV-*Tbg* viruses. ALT levels in 6-8 week old male C57BL/6J mice transduced with the indicated virus (10e11 GC/mouse) for one week. Data are shown as means ± SEM, N = 5/group.

**Supplemental Figure 3.**
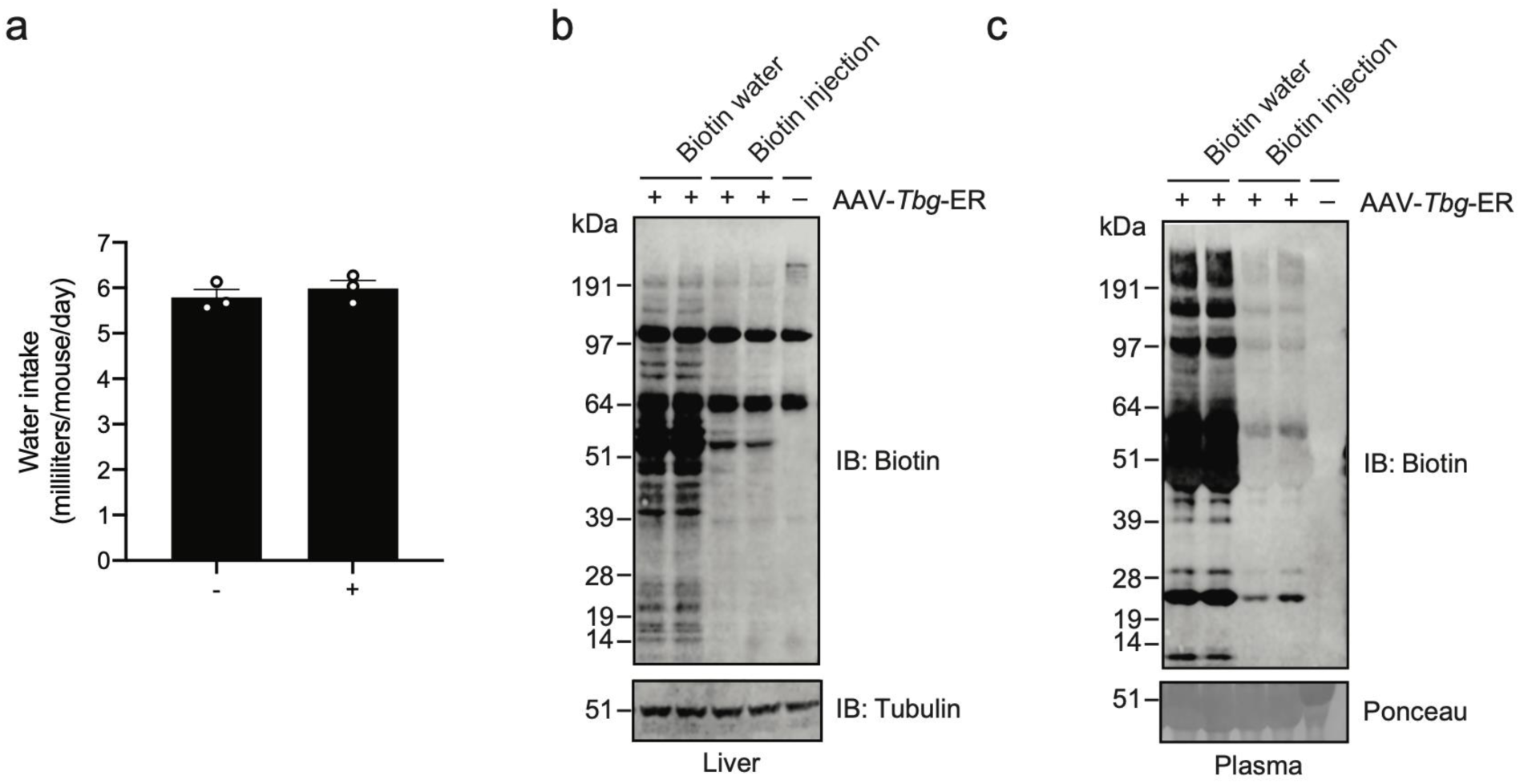
Comparison of biotin delivery methods in vivo. **(a)** Water intake of animals with regular water (left, -) or 0.5 mg/ml biotin-supplemented water (right, +). Data are shown as means ± SEM, N = 3/group. **(b, c)** Anti-biotin and anti-tubulin blotting of liver lysates (**b**) or plasma (**c**) from mice transduced with AAV-*Tbg*-ER or saline control and treated with biotin water (0.5 mg/ml for 3 days) or biotin injection (24 mg/kg/day, intraperitoneally, 3 days).

**Supplementary Figure 4.**
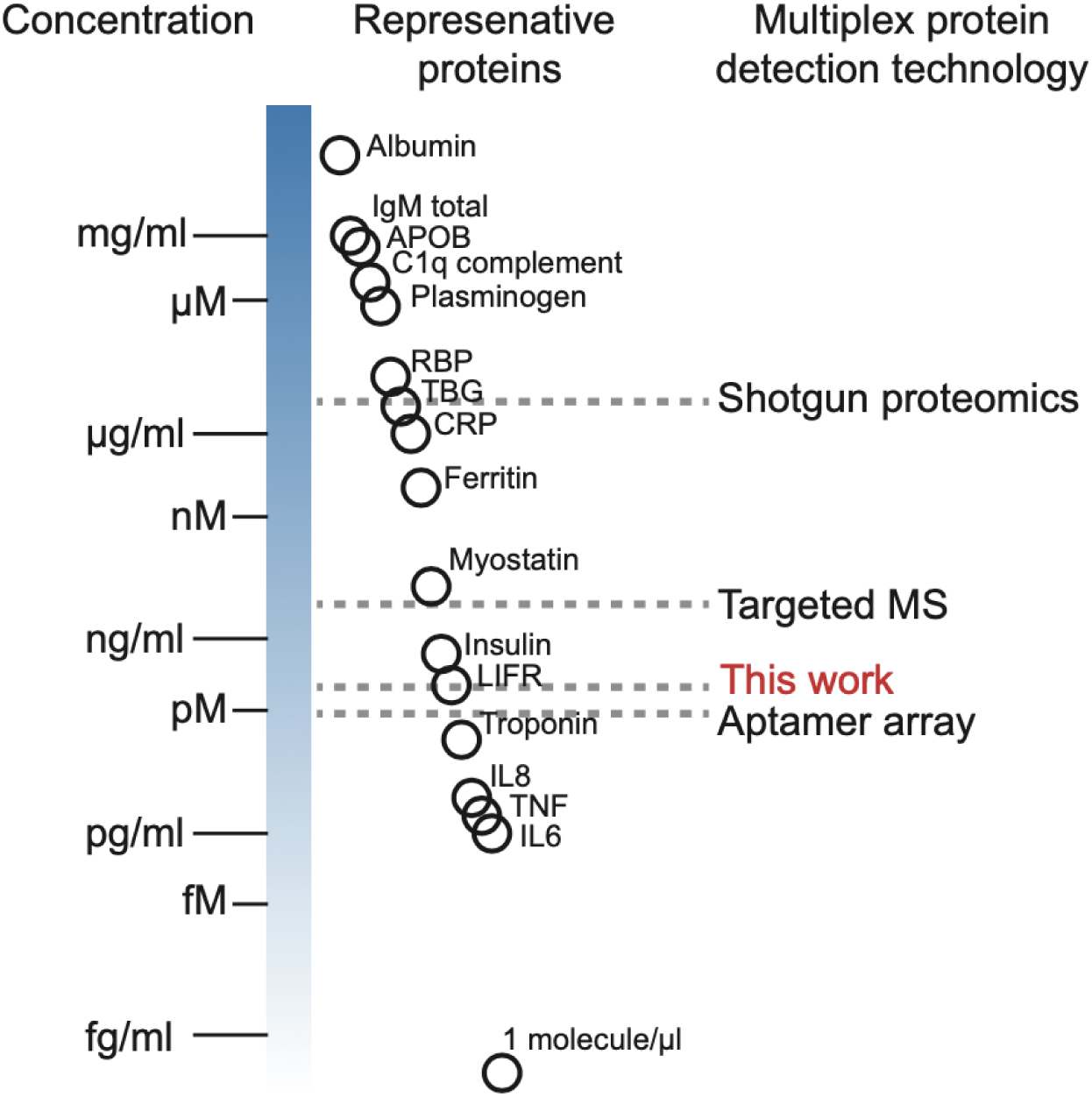
Enrichment of the hepatocyte secretome by direct purification of biotinylated proteins from blood. (**a-c**) Volcano plots of streptavidin-enriched plasma proteins from mice transduced with the hepatocyte secretome labeling reagents AAV-*Mem* (a), AAV-*Cyto* (b), or AAV-*ER* (c) versus control mice. Labeling was initiated one week after viral transduction by injection of biotin (24 mg/kg/day, 3 consecutive days). One day after the final biotin injection, plasma was harvested streptavidin-enriched, and analyzed by shotgun LC-MS/MS. N = 3mice/group.

**Supplemental Figure 5.**
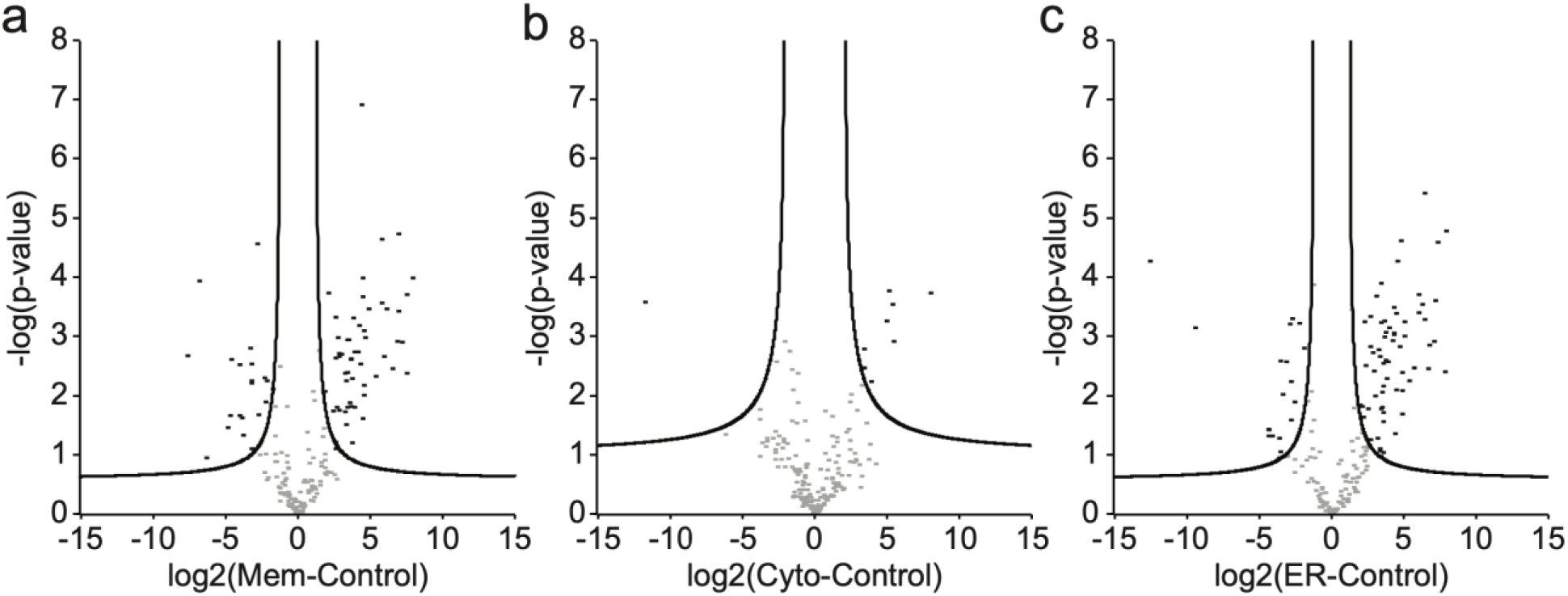
Biotinylation of intra-hepatic proteins under chow or high fructose, high sucrose diets. (**a-c**) Streptavidin blotting and anti-actin blotting of liver lysates from mice transduced with the indicated AAV-*Tbg* virus for one week, fed with chow or HFHS diet for one week and then injected with vehicle (–) or injected with biotin (24 mg/kg/day, intraperitoneally, for seven consecutive days). Livers were harvested 24 h after the final biotin injection. Quantification of total biotin signal intensity in each lane is shown to the right of the gels. Data are shown as means ± SEM, N = 3/group.

**Supplemental Figure 6.**
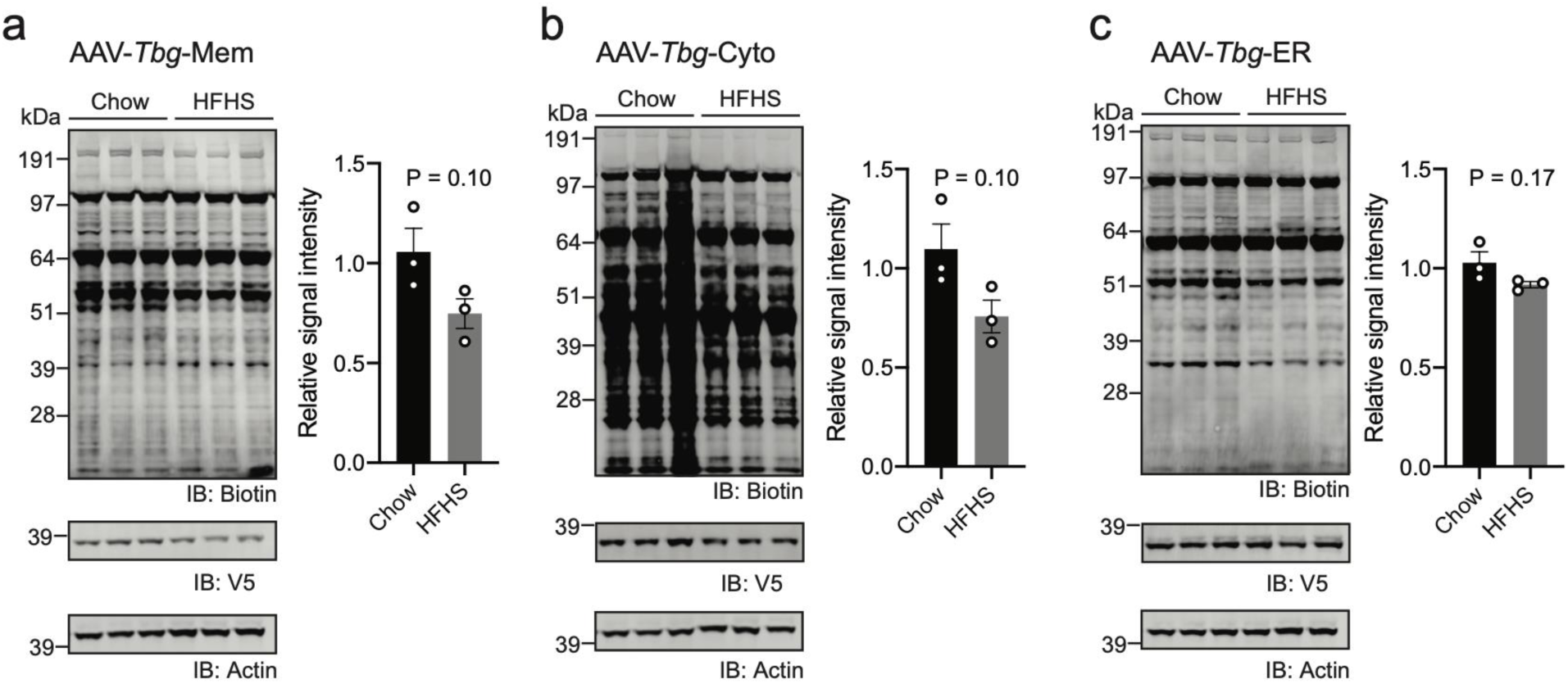
Expression of *Bhmt* mRNA across mouse tissues. *Bhmt* mRNA expression (left) and tissue legend (right) in the Tabula Muris single cell RNAseq dataset.

**Supplemental Figure 7.**
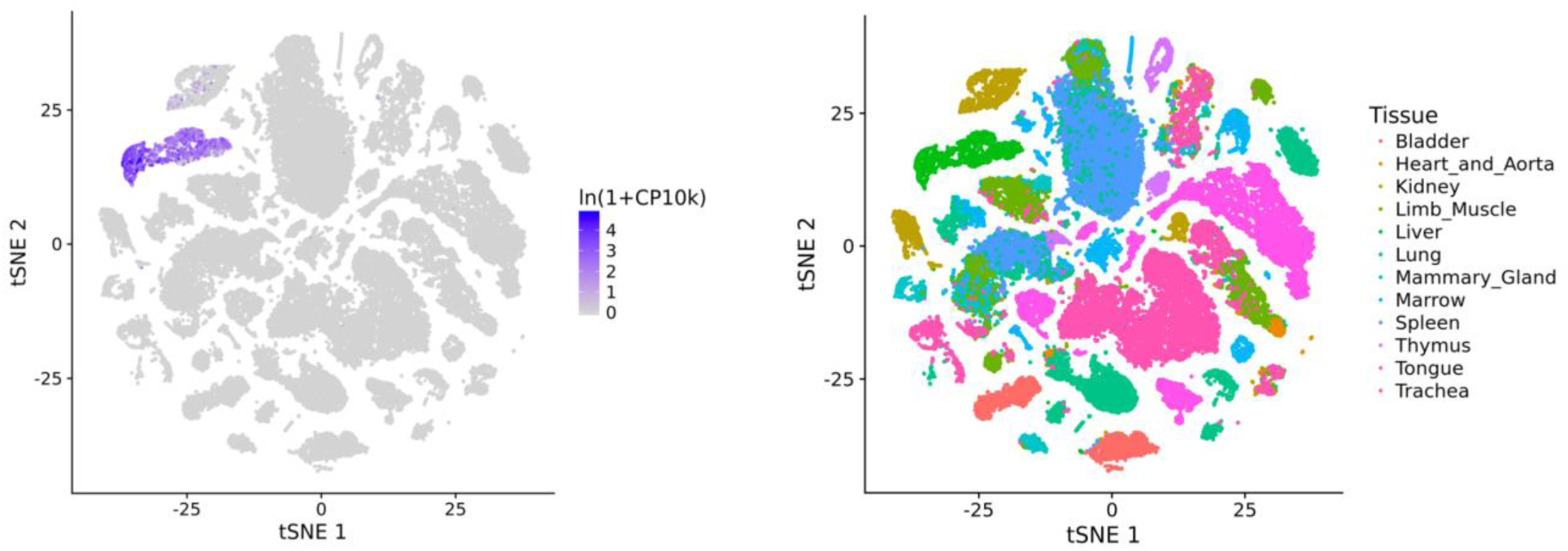
BHMT is not secreted when transfected to HEK293T cells. Anti-flag and anti-actin blotting of conditioned media and cell lysates from HEK293T cells transfected with BHMT-flag, flag-FGF2 or mock control. 24 h after transfection, cells were switched into serum-free media supplemented with 500 µM oleic acid or DMSO and incubated for additional 18 h before harvest.

**Supplemental Figure 8.**
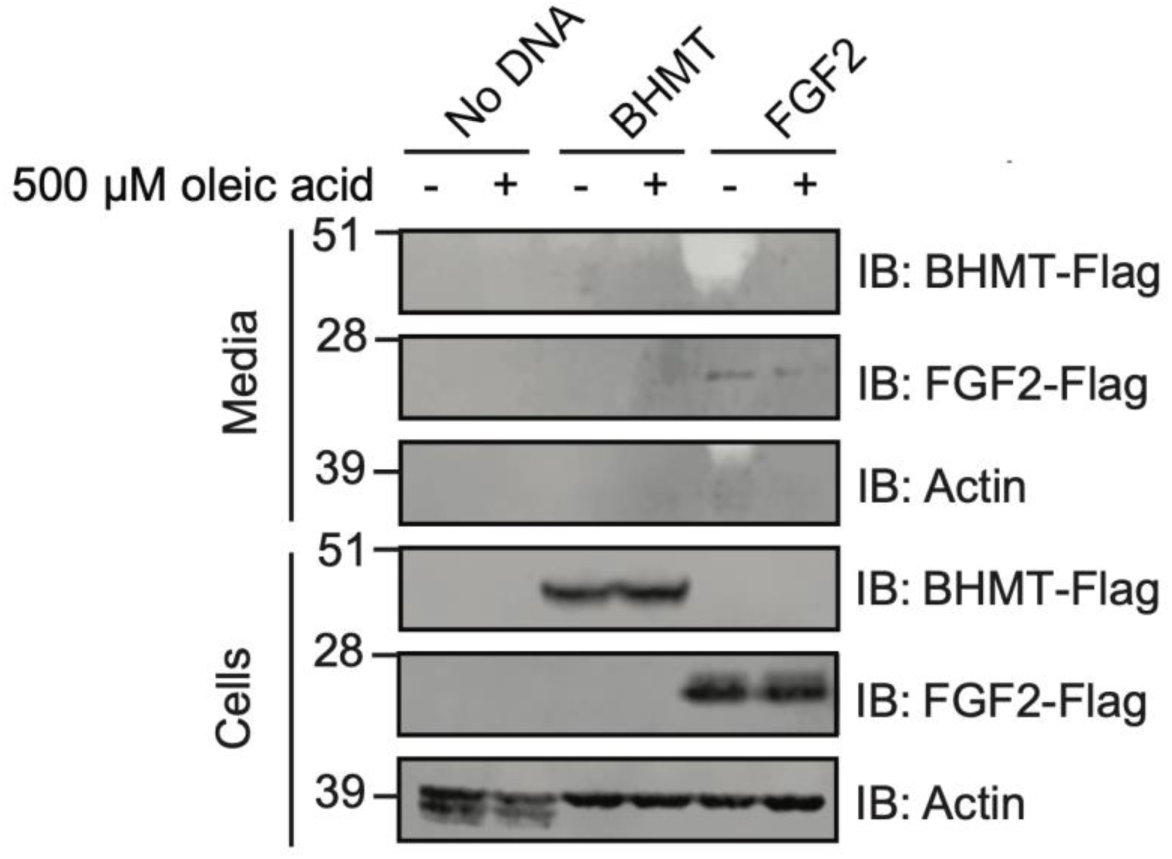
Conditional labeling strategy for myocytes, pericytes, myeloid cells, and hepatocytes in vivo. **(a)** Schematic of myocyte-specific AAV-*tMCK* virus. **(b)** Anti-V5 and anti-tubulin blotting of a panel of murine tissues following transduction by AAV-*tMCK* virus. (+) indicates AAV transduction and (–) indicates no viral transduction. Animals were C57BL/6J males age P2-3 at time of injection (10e11 GC/mouse, intraperitoneal) and tissues were harvested at 6 weeks of age. **(c)** Schematic of conditional AAV-FLEx virus. **(d)** Anti-V5, anti-biotin and anti-tubulin blotting of cell lysates from HEK293T cells transfected with the indicated plasmids. Proximity labeling was initiated three days post transfection by switching cells into serum-free media in the presence of 500 µM biotin for 18 h. Experiments were performed three times and similar results were obtained. **(e, f)** Anti-biotin blotting of liver lysates (**d**) and plasma (**e**) from 6 to 12-week old C57BL/6J male mice and *Albumin*-cre male mice transduced with AAV-FLEx virus for one week and fed with 0.5 mg/ml biotin in regular water for three consecutive days.

**Supplemental Figure 9.**
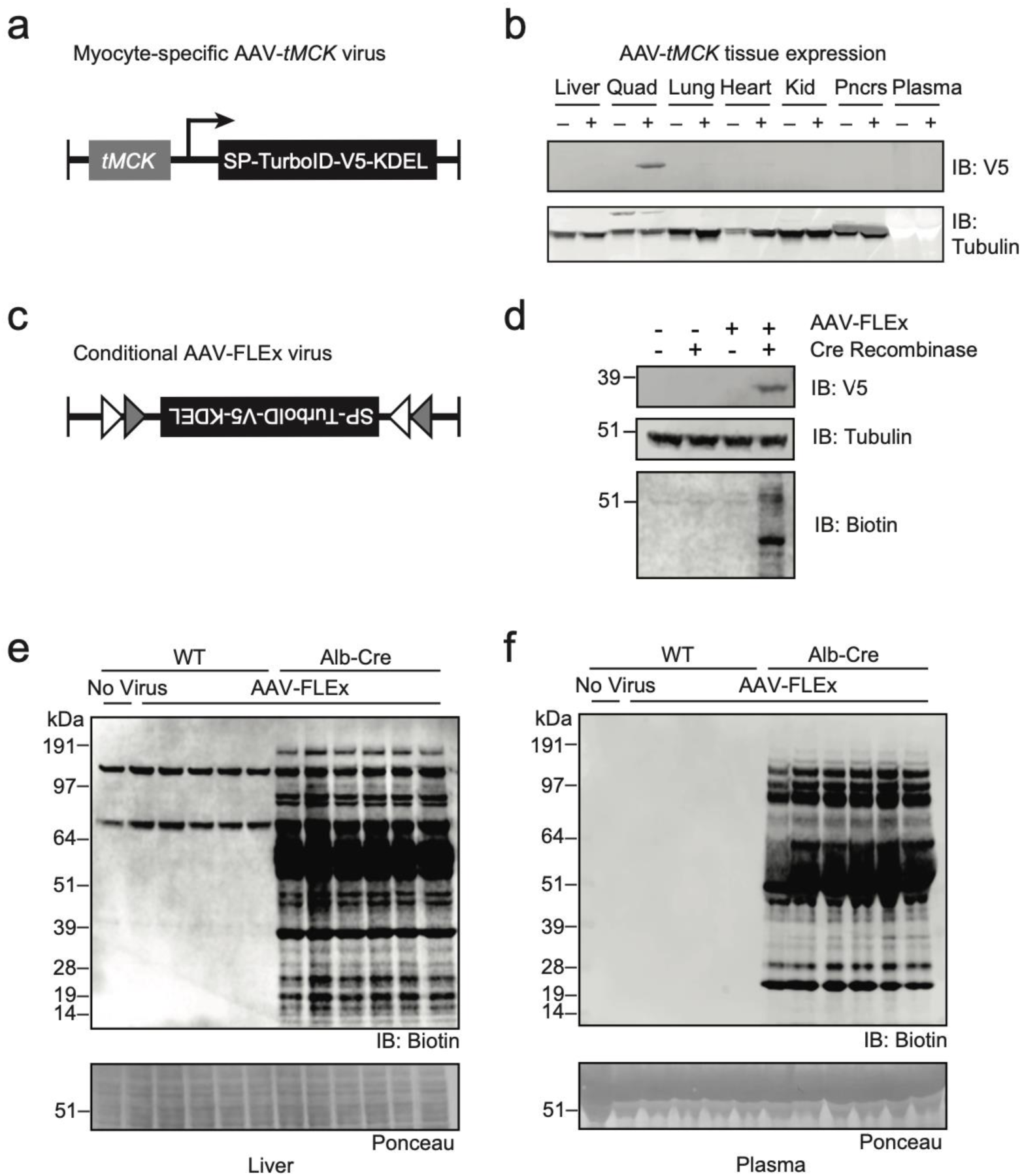
Principal component analysis of the in vivo secretomes from four different cell types. Principal component analysis of the differentially detected streptavidin-purified plasma proteins from the in vivo hepatocyte, meloid, pericyte, and myocyte secretomes.

**Supplemental Figure 10.**
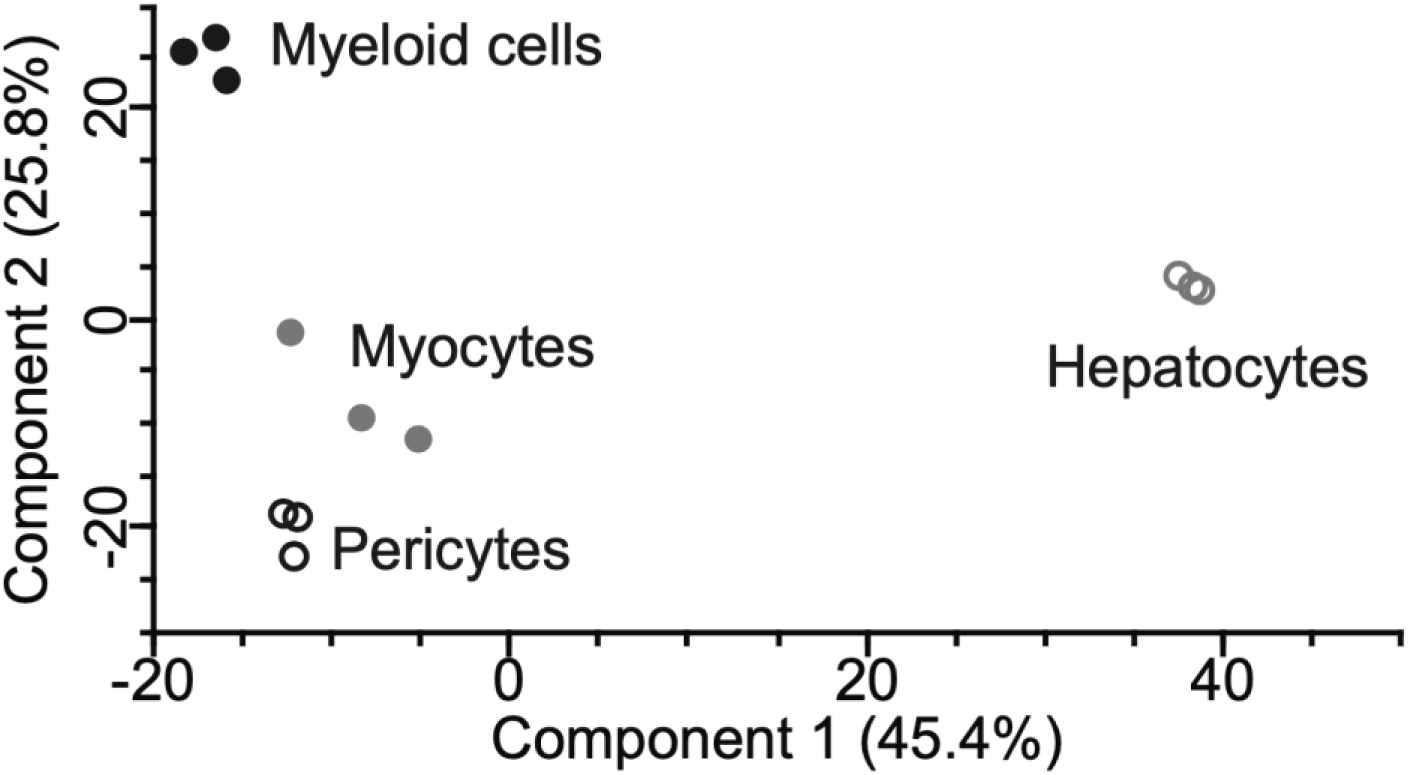
Comparison of secretome overlap between AAV-*Tbg* and *Albumin*-cre experiments. Venn diagram showing the number of proteins detected from AAV-*Tbg* and *Albumin*-cre secretomes. For the AAV-*Tbg* secretome, male 6-8 week old C57BL/6J mice were transduced with AAV-*Tbg*-ER (10e11 GC/mouse, intravenously) for one week and then injected with biotin (24 mg/kg/day, intraperitoneally, for three consecutive days). For the *Albumin-*cre secretome, AAV-FLEx virus (10e11 GC/mouse, intravenously) was injected into hemizygous 6-8 week old male *Albumin*-cre mice. After a one week transduction period, biotin was supplemented into drinking water (0.5 mg/ml, 3 days). N = 3 mice/group.

**Supplemental Figure 11.**
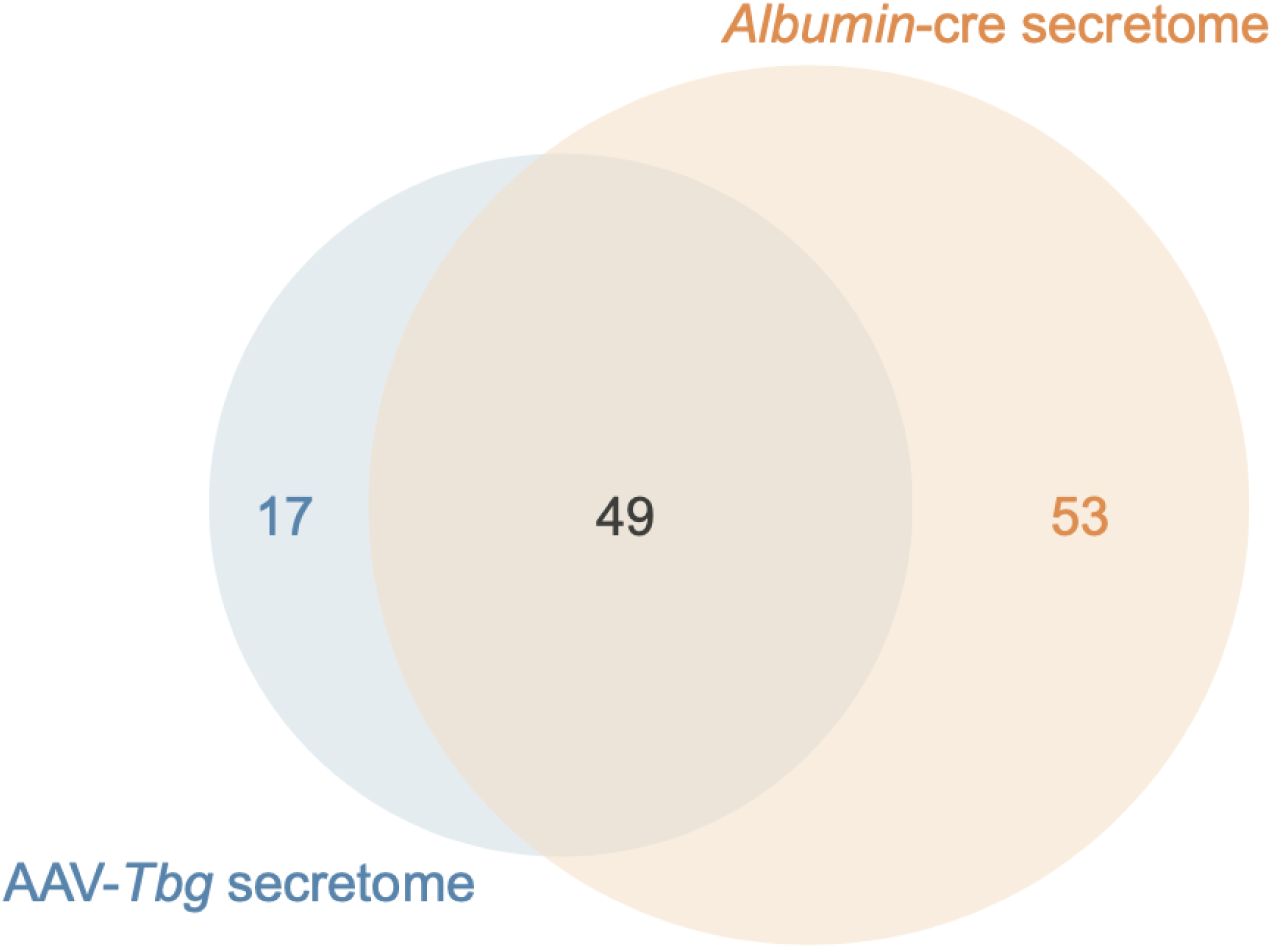
Comparison of various multiplex protein detection technologies for plasma proteomics. Absolute concentrations (left panel), representative plasma proteins (middle panel), and multiplex protein detection technology (right panel) across the entire concentration range of the plasma proteome.

**Supplemental Table 1.** LC-MS/MS proteomic analysis of streptavidin-purified plasma proteins from mice transduced with AAV-*Tbg*-Mem, AAV-*Tbg*-Cyto, or AAV-*Tbg*-ER viruses or control mice.

**Supplemental Table 2.** LC-MS/MS proteomic analysis of streptavidin-purified liver proteins from mice transduced with AAV-*Tbg*-Mem, AAV-*Tbg*-Cyto, or AAV-*Tbg*-ER viruses.

**Supplemental Table 3.** LC-MS/MS proteomic analysis of streptavidin-purified plasma from control mice (chow) without viral transduction or mice transduced with AAV-TBG-Cyto after 2 weeks of chow diet or high fructose, high sucrose diet.

**Supplemental Table 4.** LC-MS/MS proteomic analysis of streptavidin-purified plasma from mice. The following cell types correspond with the following mouse genotypes and viruses: hepatocytes (*Albumin*-cre; AAV-FLEx); myocytes (WT mice; AAV-*tMCK*); pericytes (*Pdgfrb*-creERT2; AAV-FLEx); myeloid cells (*LysM*-creERT2; AAV-FLEx).

## REFERENCES

1. Rorsman, P. & Braun, M. Regulation of insulin secretion in human pancreatic islets. Annual Review of Physiology (2013). doi:10.1146/annurev-physiol-030212-183754

2. Gaceb, A., Barbariga, M., Özen, I. & Paul, G. The pericyte secretome: Potential impact on regeneration. Biochimie (2018). doi:10.1016/j.biochi.2018.04.015

3. Lim, J. M., Wollaston-Hayden, E. E., Teo, C. F., Hausman, D. & Wells, L. Quantitative secretome and glycome of primary human adipocytes during insulin resistance. Clin. Proteomics (2014). doi:10.1186/1559-0275-11-20

4. Rabouille, C. Pathways of Unconventional Protein Secretion. Trends in Cell Biology (2017). doi:10.1016/j.tcb.2016.11.007

5. Eichelbaum, K., Winter, M., Diaz, M. B., Herzig, S. & Krijgsveld, J. Selective enrichment of newly synthesized proteins for quantitative secretome analysis. Nat. Biotechnol. (2012). doi:10.1038/nbt.2356

6. Yang, A. C. et al. Multiple Click-Selective tRNA Synthetases Expand Mammalian Cell-Specific Proteomics. J. Am. Chem. Soc. (2018). doi:10.1021/jacs.8b03074

7. Shin, J. et al. Comparative analysis of differentially secreted proteins in serum-free and serum-containing media by using BONCAT and pulsed SILAC. Sci. Rep. (2019). doi:10.1038/s41598-019-39650-z

8. Eichelbaum, K. & Krijgsveld, J. Combining pulsed SILAC labeling and click-chemistry for quantitative secretome analysis. Methods Mol. Biol. (2014). doi:10.1007/978-1-4939-0944-5_7

9. Witzke, K. E. et al. Quantitative Secretome Analysis of Activated Jurkat Cells Using Click Chemistry-Based Enrichment of Secreted Glycoproteins. J. Proteome Res. (2017). doi:10.1021/acs.jproteome.6b00575

10. Kim, D. I. et al. An improved smaller biotin ligase for BioID proximity labeling. Mol. Biol. Cell (2016). doi:10.1091/mbc.E15-12-0844

11. Roux, K. J., Kim, D. I., Raida, M. & Burke, B. A promiscuous biotin ligase fusion protein identifies proximal and interacting proteins in mammalian cells. J. Cell Biol. (2012). doi:10.1083/jcb.201112098

12. Branon, T. C. et al. Efficient proximity labeling in living cells and organisms with TurboID. Nature Biotechnology (2018). doi:10.1038/nbt.4201

13. May, D. G., Scott, K. L., Campos, A. R. & Roux, K. J. Comparative Application of BioID and TurboID for Protein-Proximity Biotinylation. Cells (2020). doi:10.3390/cells9051070

14. Octeau, J. C. et al. An Optical Neuron-Astrocyte Proximity Assay at Synaptic Distance Scales. Neuron (2018). doi:10.1016/j.neuron.2018.03.003

15. Long, J. Z. et al. The Secreted Enzyme PM20D1 Regulates Lipidated Amino Acid Uncouplers of Mitochondria. Cell (2016). doi:10.1016/j.cell.2016.05.071

16. Long, J. Z. et al. Ablation of PM20D1 reveals *N* -acyl amino acid control of metabolism and nociception. Proc. Natl. Acad. Sci. 115, 201803389 (2018).

17. Jackson, A. et al. Heat shock induces the release of fibroblast growth factor 1 from NIH 3T3 cells. Proc. Natl. Acad. Sci. U. S. A. (1992). doi:10.1073/pnas.89.22.10691

18. Yan, Z., Yan, H. & Ou, H. Human thyroxine binding globulin (TBG) promoter directs efficient and sustaining transgene expression in liver-specific pattern. Gene (2012). doi:10.1016/j.gene.2012.07.009

19. Uezu, A. et al. Identification of an elaborate complex mediating postsynaptic inhibition. Science (80-.). (2016). doi:10.1126/science.aag0821

20. Pályi-Krekk, Z. et al. EGFR and ErbB2 are functionally coupled to CD44 and regulate shedding, internalization and motogenic effect of CD44. Cancer Lett. (2008). doi:10.1016/j.canlet.2008.01.014

21. Wu, C. et al. BioGPS: An extensible and customizable portal for querying and organizing gene annotation resources. Genome Biol. (2009). doi:10.1186/gb-2009-10-11-r130

22. Samuel, V. T. Fructose induced lipogenesis: From sugar to fat to insulin resistance. Trends in Endocrinology and Metabolism (2011). doi:10.1016/j.tem.2010.10.003

23. Softic, S. et al. Divergent effects of glucose and fructose on hepatic lipogenesis and insulin signaling. J. Clin. Invest. (2017). doi:10.1172/JCI94585

24. Softic, S. et al. Dietary Sugars Alter Hepatic Fatty Acid Oxidation via Transcriptional and Post-translational Modifications of Mitochondrial Proteins. Cell Metab. (2019). doi:10.1016/j.cmet.2019.09.003

25. Schaum, N. et al. Single-cell transcriptomics of 20 mouse organs creates a Tabula Muris. Nature (2018). doi:10.1038/s41586-018-0590-4

26. Teng, Y. W., Mehedint, M. G., Garrow, T. A. & Zeisel, S. H. Deletion of betaine-homocysteine S-methyltransferase in mice perturbs choline and 1-carbon metabolism, resulting in fatty liver and hepatocellular carcinomas. J. Biol. Chem. (2011). doi:10.1074/jbc.M111.265348

27. Wang, B. et al. Construction and analysis of compact muscle-specific promoters for AAV vectors. Gene Ther. (2008). doi:10.1038/gt.2008.104

28. Schnütgen, F. et al. A directional strategy for monitoring Cre-mediated recombination at the cellular level in the mouse. Nat. Biotechnol. (2003). doi:10.1038/nbt811

29. Canli, Ö. et al. Myeloid Cell-Derived Reactive Oxygen Species Induce Epithelial Mutagenesis. Cancer Cell (2017). doi:10.1016/j.ccell.2017.11.004

30. Cuervo, H. et al. PDGFRβ-P2A-CreERT2 mice: a genetic tool to target pericytes in angiogenesis. Angiogenesis (2017). doi:10.1007/s10456-017-9570-9

31. Postic, C. et al. Dual roles for glucokinase in glucose homeostasis as determined by liver and pancreatic β cell-specific gene knock-outs using Cre recombinase. J. Biol. Chem. (1999). doi:10.1074/jbc.274.1.305

32. Zimmers, T. A. et al. Induction of cachexia in mice by systemically administered myostatin. Science (80-.). (2002). doi:10.1126/science.1069525

33. McPherron, A. C., Lawler, A. M. & Lee, S. J. Regulation of skeletal muscle mass in mice by a new TGF-β superfamily member. Nature (1997). doi:10.1038/387083a0

34. Hu, E., Liang, P. & Spiegelman, B. M. AdipoQ is a novel adipose-specific gene dysregulated in obesity. J Biol Chem 271, 10697–10703 (1996).

35. Scherer, P. E., Williams, S., Fogliano, M., Baldini, G. & Lodish, H. F. A novel serum protein similar to C1q, produced exclusively in adipocytes. J. Biol. Chem. (1995). doi:10.1074/jbc.270.45.26746

36. Delaigle, A. M., Senou, M., Guiot, Y., Many, M. C. & Brichard, S. M. Induction of adiponectin in skeletal muscle of type 2 diabetic mice: In vivo and in vitro studies. Diabetologia (2006). doi:10.1007/s00125-006-0210-y

37. Piñeiro, R. et al. Adiponectin is synthesized and secreted by human and murine cardiomyocytes. FEBS Lett. (2005). doi:10.1016/j.febslet.2005.07.098

38. Mouchiroud, M. et al. The hepatokine Tsukushi is released in response to NAFLD and impacts cholesterol homeostasis. JCI Insight (2019). doi:10.1172/jci.insight.129492

39. Barb, D., Bril, F., Kalavalapalli, S. & Cusi, K. Plasma Fibroblast Growth Factor 21 Is Associated with Severity of Nonalcoholic Steatohepatitis in Patients with Obesity and Type 2 Diabetes. J. Clin. Endocrinol. Metab. (2019). doi:10.1210/jc.2018-02414

40. Xiong, X. et al. Mapping the molecular signatures of diet-induced NASH and its regulation by the hepatokine Tsukushi. Mol. Metab. (2019). doi:10.1016/j.molmet.2018.12.004

41. Krahmer, N. et al. Organellar Proteomics and Phospho-Proteomics Reveal Subcellular Reorganization in Diet-Induced Hepatic Steatosis. Dev. Cell (2018). doi:10.1016/j.devcel.2018.09.017

42. Sun, B. B. et al. Genomic atlas of the human plasma proteome. Nature (2018). doi:10.1038/s41586-018-0175-2

43. Gold, L. et al. Aptamer-based multiplexed proteomic technology for biomarker discovery. PLoS One (2010). doi:10.1371/journal.pone.0015004

44. Hailemariam, M. et al. S-Trap, an Ultrafast Sample-Preparation Approach for Shotgun Proteomics. J. Proteome Res. (2018). doi:10.1021/acs.jproteome.8b00505

45. Tyanova, S., Temu, T. & Cox, J. The MaxQuant computational platform for mass spectrometry-based shotgun proteomics. Nat. Protoc. (2016). doi:10.1038/nprot.2016.136

46. Cox, J. et al. Andromeda: A peptide search engine integrated into the MaxQuant environment. J. Proteome Res. (2011). doi:10.1021/pr101065j

47. Cox, J. et al. Accurate proteome-wide label-free quantification by delayed normalization and maximal peptide ratio extraction, termed MaxLFQ. Mol. Cell. Proteomics (2014). doi:10.1074/mcp.M113.031591

48. Tyanova, S. et al. The Perseus computational platform for comprehensive analysis of (prote)omics data. Nature Methods (2016). doi:10.1038/nmeth.3901

49. Huang, D. W., Sherman, B. T. & Lempicki, R. A. Bioinformatics enrichment tools: Paths toward the comprehensive functional analysis of large gene lists. Nucleic Acids Res. (2009). doi:10.1093/nar/gkn923

50. Michaud, S. A. et al. Molecular phenotyping of laboratory mouse strains using 500 multiple reaction monitoring mass spectrometry plasma assays. *Commun*. Biol. (2018). doi:10.1038/s42003-018-0087-6

